# Immune landscapes predict chemotherapy resistance and immunotherapy response in acute myeloid leukemia

**DOI:** 10.1101/702001

**Authors:** Jayakumar Vadakekolathu, Mark D. Minden, Tressa Hood, Sarah E. Church, Stephen Reeder, Heidi Altmann, Amy H. Sullivan, Elena Viboch, Tasleema Patel, Narmin Ibrahimova, Sarah E. Warren, Andrea Arruda, Yan Liang, John Muth, Marc Schmitz, Alessandra Cesano, A. Graham Pockley, Peter J.M. Valk, Bob Löwenberg, Martin Bornhäuser, Sarah K. Tasian, Michael P. Rettig, Jan Davidson-Moncada, John F. DiPersio, Sergio Rutella

**Affiliations:** John van Geest Cancer Research Centre, Nottingham Trent University, Nottingham, United Kingdom; Division of Medical Oncology and Hematology, Princess Margaret Cancer Centre, Toronto, Canada; NanoString Technologies Inc., Seattle, WA, United States of America; Department of Medicine, Universitätsklinikum Carl Gustav Carus, Dresden, Germany; Department of Pediatrics, Division of Oncology and Centre for Childhood Cancer Research, Children’s Hospital of Philadelphia and University of Pennsylvania School of Medicine, PA, United States of America; MacroGenics Inc., Rockville, MD, United States of America; Institute of Immunology, Faculty of Medicine Carl Gustav Carus, Technische Universität Dresden, Dresden, Germany; National Center for Tumor Diseases (NCT), Partner Site Dresden, Dresden, Germany; German Cancer Consortium (DKTK), Partner Site Dresden, and German Cancer Research Center (DKFZ), Heidelberg, Germany; Centre for Health, Ageing and Understanding Disease (CHAUD), Nottingham Trent University, Nottingham, United Kingdom; Department of Hematology, Erasmus University Medical Centre, Rotterdam, Netherlands; Division of Oncology, Department of Internal Medicine, Washington University in St. Louis, St. Louis, MO, United States of America

## Abstract

This study dissected the complexity of the immune architecture of acute myeloid leukemia (AML) at high resolution and assessed its influence on therapeutic response. Using 387 primary bone marrow samples from three discovery cohorts of children and adults with AML, we defined immune-infiltrated and immune-depleted disease subtypes and unraveled critical differences in immune gene expression across age groups and disease stages. Importantly, interferon (IFN)-γ-related mRNA profiles were predictive for both chemotherapy resistance and response of primary refractory/relapsed AML to flotetuzumab immunotherapy. Our compendium of microenvironmental gene and protein profiles sheds novel insights into the immuno-biology of AML and will inform the delivery of personalized immunotherapies to IFN-γ-dominant AML subtypes.

---

Acute myeloid leukemia (AML) is a molecularly and clinically heterogeneous hematological malignancy^1^. The discovery of the genomic landscape of AML, including the identification of targetable mutations^2^, has propelled the development of novel anti-leukemic agents and is enabling disease classification and patient stratification into favorable, intermediate or adverse risk groups^3^. Despite success in many areas, AML is cured in only 35-40% of patients <60 years of age and in 5-15% of patients >60 years of age. While chemotherapy resistance is common, the majority of patients die of disease relapse. Investigation of new molecularly-targeted and immuno-modulatory agents therefore remains a high priority for both children and adults^4^.

Tumor phenotypes are dictated not only by the neoplastic cell component, but also by the immunologic milieu within the tumor microenvironment (TME), which is equipped to subvert host immune responses and hamper effector T-cell function^5^. *In silico* approaches have been instrumental for the identification of immunogenomic features with therapeutic and prognostic implications. In solid tumors, six immune subtypes have been described (wound healing, interferon (IFN)-γ-dominant, inflammatory, lymphocyte-depleted, immunologically quiet, and transforming growth factor (TGF)-β-dominant). These are characterized by differences in macrophage or lymphocyte signatures, T helper type (Th)-1 to Th2 cell ratio, extent of intratumoral heterogeneity and neoantigen load, aneuploidy, cell proliferation, expression of immunomodulatory genes, and patient survival^6^.

Although immunotherapy may be an attractive modality to exploit in patients with AML^7^, the ability to predict the groups of patients and the forms of leukemia that will respond to immune targeting remains limited^8–11^. Clinical studies in patients with solid tumors have shown that responses to anti-programmed cell death 1 (PD-1)/programmed death ligand 1 (PD-L1)-targeted immunotherapy occur most often in individuals with immune-inflamed lesions that are characterized by pre-existing CD8^+^ T-cell responses, release of pro-inflammatory and effector cytokines^12–14^, and an augmented T-cell receptor (TCR) clonal diversity pre-treatment^15,16^. The above T-cell inflamed gene expression profile (GEP) is a measure of IFN-γ-responsive genes that are related to adaptive immune resistance (AIR) mechanisms of immune escape such as indoleamine 2,3-dioxygenase-1 (*IDO1*) and *PD-L1*^17^, and is predictive of clinical benefit with pembrolizumab immunotherapy^18,19^. Although IFN-γ plays a critical role in eliciting anti-tumor T-cell activity and enabling tumor rejection, prolonged IFN-γ signaling under conditions of persistent antigen exposure has been shown to activate a PD-L1-independent, STAT1-driven multigenic program which confers resistance to radiotherapy and anti-cytotoxic T-lymphocyte antigen 4 (CTLA-4) immunotherapy in mouse models of melanoma^20^.

Herein, we used targeted immune gene expression profiling (IGEP) and spatially-resolved multiplexed digital spatial profiling (DSP) for the high-dimensional analysis of the immunological contexture of a broad collection of bone marrow (BM) samples from patients with AML and for the identification of molecular determinants of immunotherapeutic benefit. We reveal unifying immune features and critical differences that define classes and subclasses of TMEs and deliver predictions of chemotherapy resistance, survival and immunotherapy response that are beyond the current capabilities of single molecular markers.

## Results

### Targeted immune gene expression profiling (IGEP) identifies immune subtypes of AML

We first analyzed unfractionated, archival BM samples from treatment-naïve patients with non-promyelocytic AML (PMCC discovery series; n=290 cases; **Table 1**)^21^. We derived immune scores from mRNA expression levels, similar to those of previous publications, and devised an RNA-based, quantitative metric of immune infiltration^22,23^. As shown in **Extended Data Fig. 1A-B**, patients with adverse cytogenetic features exhibited a shorter relapse-free survival (RFS) and overall survival (OS) compared with patients with intermediate and favorable cytogenetic risk, thus confirming the overall trends of well-established European Leukemia-Net (ELN) categories^24^. A Pearson correlation matrix of immune gene sets allowed us to identify co-expression patterns of pre-defined immune cell types and immune biological activities. Immune signature modules in pre-treatment BM samples reflected the co-expression of genes associated with 1) IFN-γ biology, 2) adaptive immune responses, and 3) myeloid cell abundance (macrophages, neutrophils and dendritic cells (DCs); **Fig. 1A**). We then computed the sum of the individual scores in each signature module in **Fig. 1A** and generated three immune scores (IFN-dominant, adaptive and myeloid) which individually separated AML cases according to high and low expression values (**Extended Data Fig. 1C**). When considered in aggregate, IFN-dominant, adaptive and myeloid scores dichotomized BM samples into two immune subtypes, which will be herein termed immune-infiltrated and immune-depleted (**Fig. 1B**)^25^. These immune subtypes of AML expressed comparable levels of leukemia-associated antigens CD34, CD123 (*IL3RA*) and CD117 (*KIT*), suggesting that targeted IGEP of bulk BM specimens largely captured elements of the immunological TME rather than features of the tumor cell compartment (**Extended Data Fig. 1D**).

**Table 1.** AML cohorts selected for targeted immune gene expression profiling

| Patient series | PMCC | CHOP | SAL |
| --- | --- | --- | --- |
| # of patients | 290 | 39 | 38 |
| Age (0-39), n | 76 | 39 | 11 |
| Age (40-59), n | 126 | 0 | 15 |
| Age (>60), n | 88 | 0 | 12 |
| Median age (years, range) | 52 (18-81) | 10 (0.1-20) | 52.5 (23-75) |
| WBC count at presentation | $19.15 \times 10^3/\mu\text{L}$<br>(0.7-399) | N.A. | $60.75 \times 10^3/\mu\text{L}$<br>(0.84-320.2) |
| Cytogenetic risk group, n |  |  |  |
| ELN favorable | 35 | 8 | 7 |
| ELN intermediate | 155 | 24 | 27 |
| ELN adverse | 59 | 7 | 2 |
| N.A. | 41 | - | 2 |
| Disease status at time of BM sampling, n |  |  |  |
| Diagnosis | 290 | 39 | 38 |
| Complete remission | 0 | 0 | 11 |
| Relapse | 0 | 0 | 9 |
| # of BM samples analyzed | 290 | 39 | 58 |
| # of BM samples analyzed using the GeoMx<sup>®</sup> DSP platform | 0 | 0 | 10 |
| *De novo*/secondary/therapy-related | 244/46/0 | 38/1/0 | 31/5/2 |
| Response to induction chemotherapy |  |  |  |
| Yes | 210 | 29 (M1*) | 33 |
| No | 80 | 4 (M2/M3*) | 2 |
| N.A. | - | 6 | 3 |
| Relapse |  |  |  |
| Yes | 132 | 24 | 17 |
| No | 118 | 13 | 18 |
| Induction failure | 39 | 2 | 0 |
| N.A. | 1 | - | 3 |
| Median follow-up time (months) | 101.23 | 39.9 | 16.89 |
| Median relapse-free survival (months) | 19.1 | 14.7 | 18.3 |
| Median overall survival (months) | 21.37 | 44.8 | 25.67 |

**Legend:** N.A. = Not available; PMCC = Princess Margaret Cancer Centre; CHOP =Children's Hospital of Philadelphia; SAL = Studienallianz Leukämie; ELN = European Leukemia Net; BM = bone marrow; DSP = digital spatial profiling; N.A. = not available. Median follow-up time was calculated using the reverse Kaplan-Meier method, with the event indicator reversed so that the outcome of interest becomes censored. \*M1, M2 and M3 BM remission status was defined as <5%, 5% to 24% and >25% AML blasts after induction chemotherapy, respectively<sup>63</sup>.

**Fig. 1:**
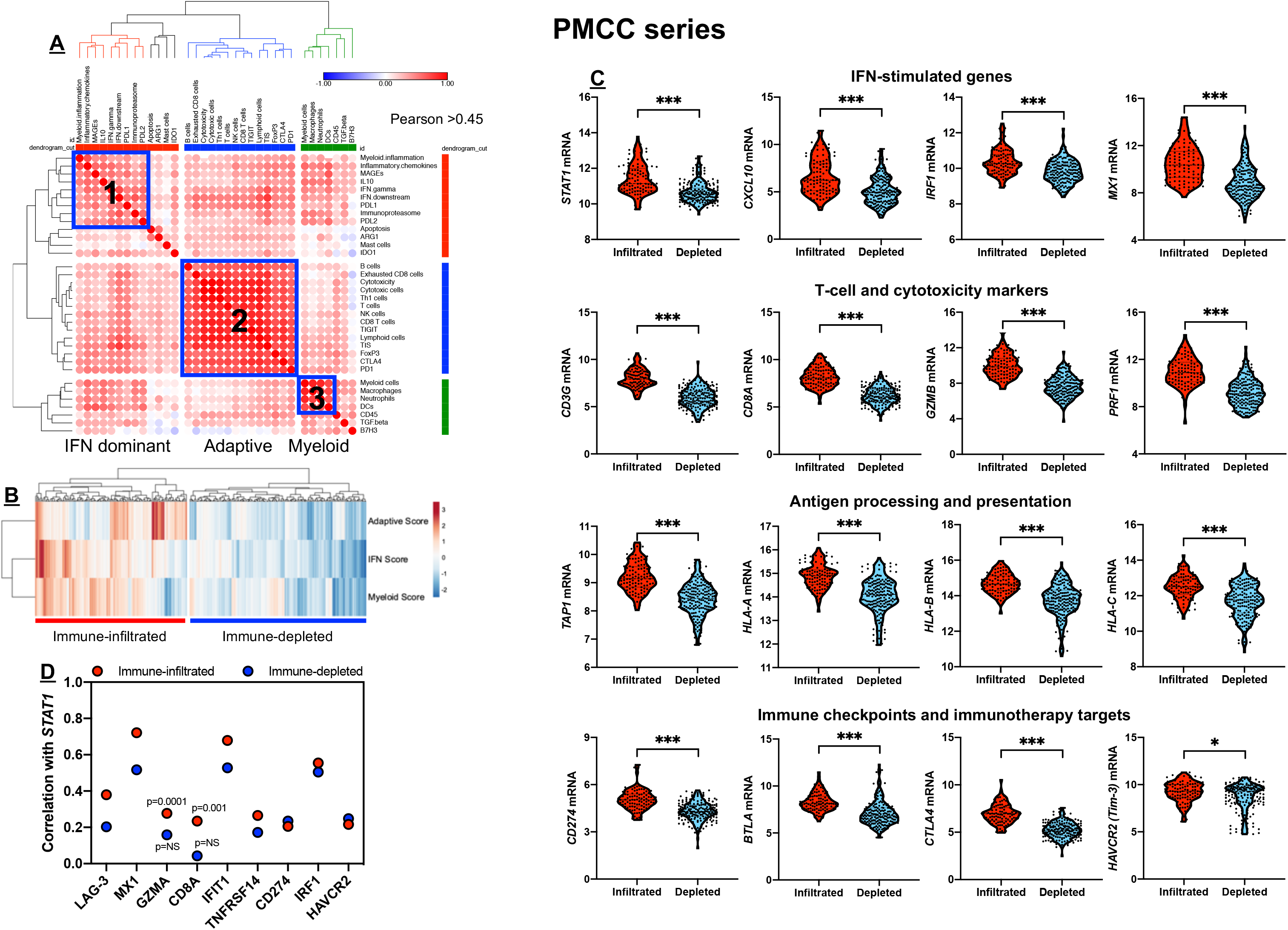
Immune gene sets stratify bone marrow samples from patients with newly diagnosed AML (PMCC cohort). **A**) Unsupervised hierarchical clustering (Euclidean distance, complete linkage) of the correlation matrix of immune and biological activity signatures identifies co-expression patterns (blue boxes) of immune gene sets (correlation value color-coded *per* the legend; Pearson correlation coefficient >0.45) in the bone marrow (BM) microenvironment of patients with AML, namely, IFN-dominant (1), adaptive (2) and myeloid (3) gene modules. Immune cell type^23^ and signature scores^22^ were calculated from mRNA levels as pre-defined linear combinations (weighted averages) of biologically relevant gene sets. Morpheus, an online tool developed at the Broad Institute (MA, USA) was used for data analysis and visualization. **B**) IFN-dominant, adaptive and myeloid scores in aggregate stratify patients with newly diagnosed AML into two distinct clusters, which are referred in this study as immune-infiltrated and immune-depleted^25^. ClustVis, an online tool for clustering of multivariate data, was used for data analysis and visualization^64^. **C**) Violin plots summarizing the expression of IFN-stimulated genes (ISGs), T-cell and cytotoxicity markers, negative immune checkpoints, genes implicated in antigen processing and presentation, and immunotherapy targets in AML cases with an immune-infiltrated and immune-depleted tumor microenvironment (TME). Data were compared with the Mann-Whitney *U* test for unpaired determinations (two-sided). *P<0.05; ***p<0.0001. D) Correlation between *STAT1*, ISGs [IRF1, *MX1, IFIT1, TNFRSF14, PD-L1* (CD274)], surrogate markers for cytotoxic T cells (*CD8A, GZMA*) and negative immune checkpoints [*LAG3, HAVCR2* (Tim-3)] under conditions of high and low immune infiltration.

As shown in **Fig. 1C**, AML cases with immune-infiltrated profiles expressed significantly higher levels of IFN-stimulated genes and T-cell recruiting factors (*STAT1, CXCL10, IRF1*), T-cell markers and cytolytic effectors (*CD8A, CD8B, GZMB, PRF1*), counter-regulatory immune checkpoints and immunotherapy drug targets (*IDO1, CTLA4, PD-L1* and *BTLA*), and molecules involved in antigen processing and presentation (*TAP1, TAP2, HLA-A, HLA-B* and *HLA-C*). Conceivably, high T-cell infiltration, MHC expression and PD-L1 levels in the immune-infiltrated AML subtype reflected a pre-existing IFN-γ-driven adaptive immune response which has previously been associated with suppressed anti-tumor immune reactivity^13,26^, but also with immunotherapy responses in patients with solid tumors^9,18,27^ and AML^11^. The expression of *STAT1*, a central component of the IFN-γ signaling pathway and predictor of response to immune checkpoint blockade^28^, was more strongly correlated with the presence of T-cell inhibitory receptors *TNFRSF14* (a ligand for the immunoglobulin superfamily members BTLA and CD160), *PD-L1, HAVCR2* (Tim-3) and *LAG-3*, and with IFN-stimulated genes *MX1, IFIT1* and *IRF1* in the immune-infiltrated relative to the immune-depleted subtype, consistent with their coordinated regulation in an inflamed TME (**Fig. 1D**). Finally, *PD-L1, IFN-γ* signaling, IFN downstream signaling and immunoproteasome mRNA scores in the IFN-dominant gene module, but not adaptive and myeloid gene modules (data not shown), individually separated patients into subgroups with different survival probabilities (**Extended Data Fig. 1E**).

### Highly multiplexed digital spatial profiling (DSP) unravels distinct T-cell neighbors in immune-infiltrated and immune-depleted AML

Transcriptomic data do not provide information on spatial relationships of tumor-infiltrating immune cells within the TME. Therefore, we used GeoMx^®^ DSP to characterize the expression of 31 immuno-oncology (IO) proteins in 10 fresh frozen paraffin embedded (FFPE) BM biopsies from treatment-naïve patients with AML (SAL patient series) with varying degrees of T-cell infiltration (**Extended Data Fig. 2A**). We selected 24 geometric regions of interest (ROIs) per BM sample using fluorescent anti-CD3 (visualization marker for T cell-rich ROIs) and anti-CD123 antibodies (visualization marker for myeloid blast-rich ROIs)^29^ (**Extended Data Fig. 3–4**). T-cell gene expression scores (calculated as detailed in **Extended Data Fig. 2B**) correlated with DSP CD3 protein status (**Fig. 2A** and **Extended Data Fig. 2C**). Furthermore, mRNA and protein levels for B cells, monocytes and Bcl-2 significantly and positively correlated, serving as a validation for the mRNA-based immune scores (**Extended Data Fig. 2D**).

**Fig. 2:**
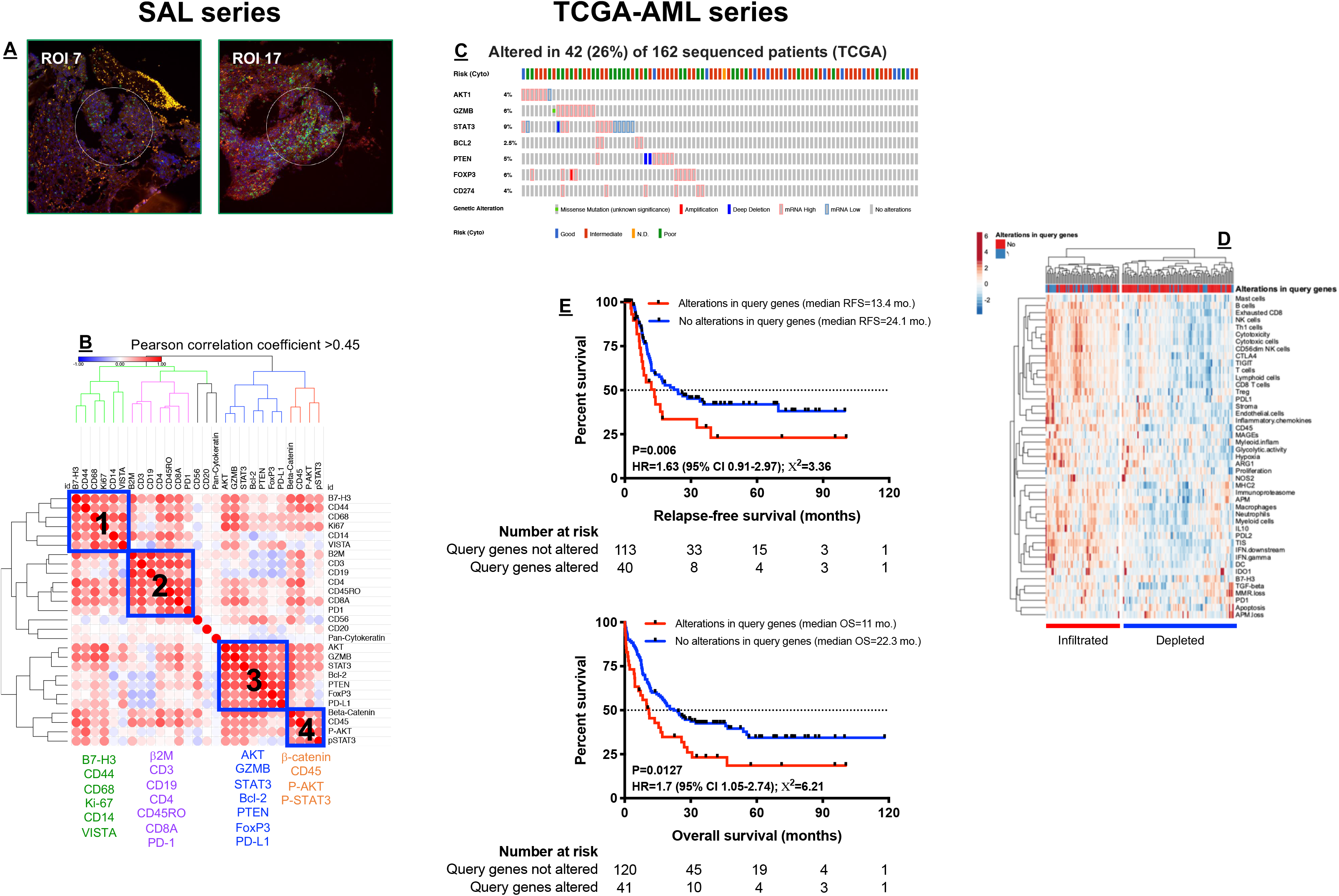
Multiplexed protein detection with GeoMx™ Digital Spatial Profiling (DSP) identifies prognostic signatures in immune-infiltrated AML. **A**) CD3 expression (green fluorescence) in representative regions of interest (ROIs) from a bone marrow (BM) trephine biopsy obtained at time of AML diagnosis (SAL series). **B**) Correlation matrix of protein expression in BM biopsies from 10 patients with newly diagnosed AML (SAL series). Protein expression data were subjected to unsupervised hierarchical clustering. Heat-maps were built using Morpheus (Broad Institute, MA) with blue boxes denoting four distinct protein coexpression patterns (Pearson correlation coefficient >0.45) or signatures (SIG). **C**) Abnormalities in SIG3 proteins were detected in 26% of TCGA-AML cases (42 of 162 sequenced BM samples). Data were retrieved, analyzed and visualized using cBioPortal. **D**) Heat-map of immune cell type-specific scores and biological activity scores in TCGA-AML cases with and without abnormalities of SIG3 genes. ClustVis, an online tool for clustering of multivariate data, was used for data analysis and visualization^64^. **E**) Kaplan-Meier estimates of relapse-free survival (RFS) and overall survival (OS) in TCGA-AML patients with (red line) and without (blue line) abnormalities of SIG3 genes. HR = hazard ratio. Survival curves were compared using a log-rank (Mantel-Cox) test.

We then asked whether CD3-rich and CD3-poor BM samples (defined by a median split of barcode counts) differed in terms of co-localization patterns of relevant IO proteins. As shown in **Extended Data Fig. 2A**, CD3^+^ T cells in immune-infiltrated biopsies co-localized with B cells, antigen processing and presentation-related proteins (β2-microglobulin), negative immune checkpoint B7-H3 and β-catenin. In contrast, CD3^+^ T cells in immune-depleted biopsies co-localized with markers of immunological memory (CD45RO) and T-cell exhaustion (PD-1). When analyzing overall protein expression patterns (10 samples × 24 ROIs *per* sample × 31 proteins = 7,440 data points), we identified four protein signatures (SIG), which were then further assessed *in silico* for correlations with clinical-biological disease characteristics and potential prognostic value in The Cancer Genome Atlas (TCGA)-AML cases (162 sequenced AML samples with putative copy-number alterations, mutations and mRNA expression z-scores [threshold±2.0]). Abnormalities in SIG1, SIG2 and SIG4 genes (blue boxes in **Fig. 2B**) did not correlate with specific disease characteristics or clinical outcomes (data not shown). In contrast, mRNA up-regulation, gene amplification, deep deletion and mis-sense mutations in SIG3 genes, which were detected in 26% of TCGA-AML cases (**Fig. 2C**), significantly correlated with *TP53* mutation status, an established adverse prognosticator in AML (*p* value from mutation enrichment analysis = 0.0285). Alterations in SIG3 genes, which included *PD-L1, FoxP3*, molecules associated with cell cytotoxicity, *PTEN* and *BCL2*, were predominantly observed in patients with immune-infiltrated mRNA profiles (**Fig. 2D**) and also correlated with higher number of mutations (median=12 and 9 in AML cases with [n=41] or without [n=115] abnormalities in SIG3 genes, respectively; p=0.021) and with adverse ELN cytogenetic features (χ^2^=25.03; p<0.001), but not with other disease characteristics at presentation, including white blood cell (WBC) count, and percentage of AML blasts in pre-treatment blood and BM samples (data not shown). Finally, patients with abnormalities in SIG3 genes experienced poor clinical outcomes, as shown by the significantly lower RFS and OS rates (**Fig. 2E**). Collectively, highly multiplexed *in situ* detection of IO proteins highlights critical differences in T-cell infiltrated *versus* T-cell depleted AML subtypes and identifies protein signatures with prognostic potential in T-cell infiltrated pre-treatment samples.

### Interactions between immune subgroups, common cytogenetic alterations and clinical factors

We next correlated immune signature scores with clinical and demographic factors, including WBC count and blast cell count at diagnosis, ELN cytogenetic category (available in 249 cases from the PMCC discovery series) and patient age. Leukemia burden was significantly lower in the immune-infiltrated AML subtype (median WBC count at diagnosis=10.0×10^3^/μL, range 1.0-188.0, *versus* 25.8×10^3^/μL, range 0.7-398.3, p<0.0001; median percentage of BM blasts=50% *versus* 80%, p<0.0001, and median number of AML blasts per μL of blood=3.7 *versus* 10.71, p<0.0001; **Extended Data Fig. 5A-C**). Although immune signature scores were not correlated with the ELN cytogenetic risk category when considered individually (data not shown), we observed a trend towards more AML cases with ELN adverse cytogenetics in the immune-infiltrated subtype (32% *versus* 20%, p=0.021; χ^2^ test for trend; **Extended Data Fig. 5D**). Finally, patients with high immune infiltration tended to be of a more advanced age at diagnosis (median=58 years, range 23-79) compared with patients with low immune infiltration (median=50 years, range 18-63, p<0.0001; **Extended Data Fig. 5E**).

### Immune subtypes improve survival prediction

The activation of immune pathways has context-dependent prognostic impact that differs between tumor types^6^. We first assessed the ability of the immune subtype to refine the accuracy of outcome prediction separately for each ELN cytogenetic risk category. Among patients with favorable risk, RFS and OS times were significantly longer in individuals with an immune-infiltrated TME (**Fig. 3A**). In contrast, clinical outcomes in ELN adverse risk cases were worse in individuals with an immune-infiltrated TME. We also observed that the ELN classifier assisted outcome prediction only in the immune-infiltrated subtype (**Fig. 3B**), allowing the identification of patient subgroups with excellent survival estimates (87.5% RFS and 77.8% OS) or with very unsatisfactory outcomes [10.4% RFS (log-rank χ^2^=15.07; p<0.0001) and 7.2% OS (log-rank χ^2^=25.75; p<0.0001); **Fig. 3B**]. This finding is congruent with previous studies showing that specific gene expression profiles correlate with longer OS in patients with favorable-risk AML, but not in those with intermediate-risk or high-risk AML^30^. Similarly, leukemia stem cell-associated gene signatures have been shown to provide prognostic separation within specific ELN risk categories^31^.

**Fig. 3:**
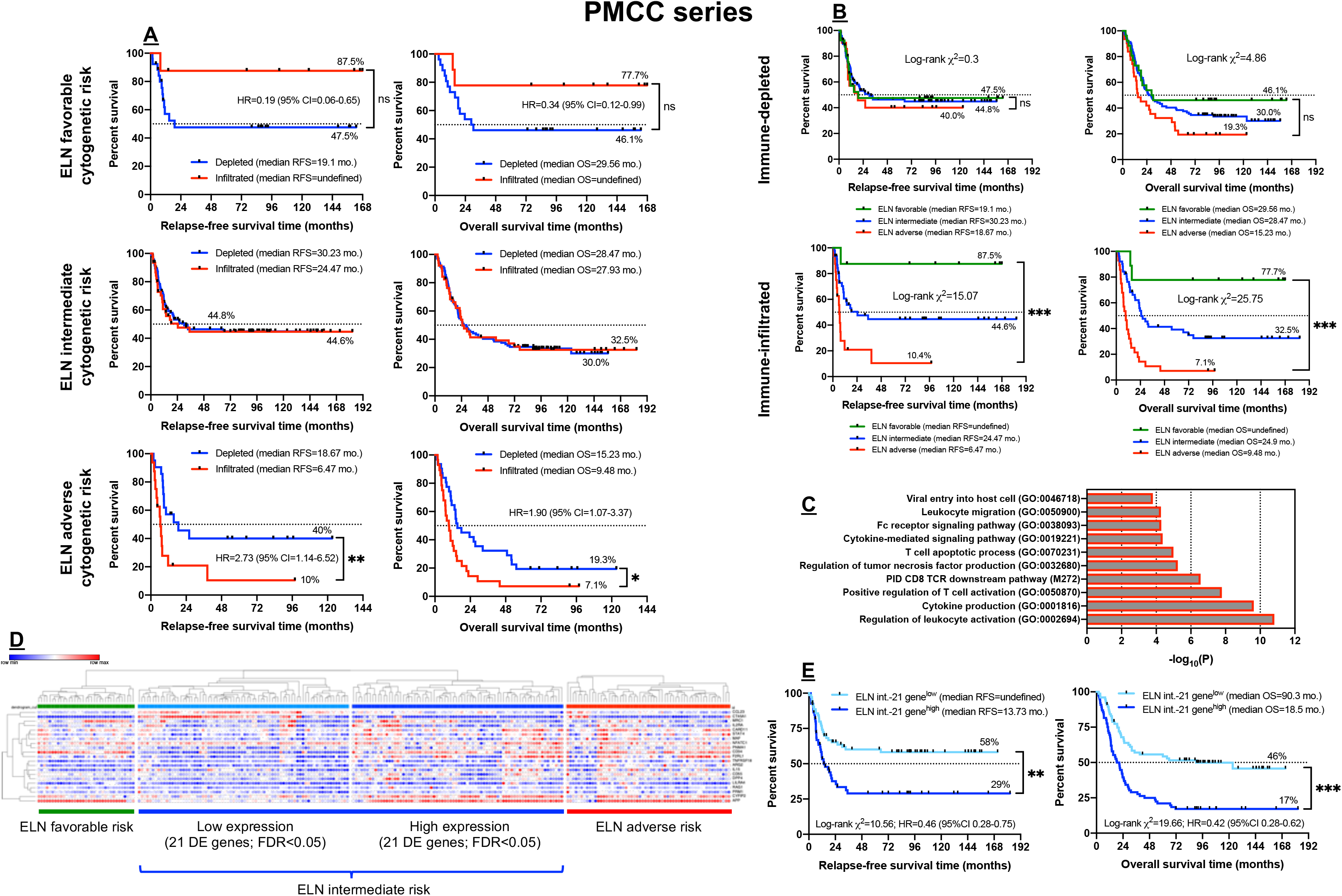
Clinical correlates of immune profiles in patients with newly diagnosed AML (PMCC cohort). **A**) Stratification of patient survival within each ELN cytogenetic risk category by immune subtype (immune-infiltrated and immune-depleted). Kaplan-Meier estimates of RFS and OS are shown. Survival curves were compared using a log-rank (Mantel-Cox) test. *P<0.05; **P<0.01. HR=hazard ratio; CI=confidence interval. **B**) Cytogenetically-defined categories stratify survival in patients with immune-infiltrated AML. Kaplan-Meier estimates of RFS and OS are shown. Survival curves were compared using a log-rank (Mantel-Cox) test. ***P<0.001. **C**) Differentially expressed (DE) genes (n=21) between favorable and adverse-risk AML were mapped to gene ontology (GO) biological processes and pathways using Metascape.org. **D**) Expression of the 21 DE genes across the PMCC discovery cohort (unsupervised hierarchical clustering; Euclidean distance; complete linkage). FDR=false discovery rate. Morpheus, an online tool developed at the Broad Institute (MA, USA), was used for data analysis and visualization. **E**) Kaplan-Meier estimate of RFS and OS in patients with ELN intermediate-risk AML stratified by the 21-gene classifier. Survival curves were compared using a log-rank (Mantel-Cox) test. HR=hazard ratio; CI=confidence interval. **P<0.01; ***P<0.001.

Unexpectedly, our immunological classifier was unable to stratify survival in patients with intermediate ELN risk (**Fig. 3A**). We therefore employed an immune gene signature-agnostic approach to identify gene sets with prognostic impact in this specific patient category. By performing Cox proportional hazards regression, we discovered a set of 21 differentially expressed (DE) immune genes [false discovery rate (FDR)<0.05] between favorable and adverse-risk AML (**Supplemental Table 1**), which were significantly associated with OS and exhibited enrichment of gene ontologies (GO) and pathways related to T-cell activation, TCR downstream signaling and regulation of cytokine production (**Fig. 3C**). Interestingly, patients with intermediate-risk AML could be separated into subgroups with low and high gene expression values, with the former being closely similar to ELN favorable-risk patients while the latter resembled ELN adverse-risk patients (**Fig. 3D**). Importantly, RFS and OS estimates were significantly poorer for intermediate-risk patients with high *versus* low expression levels of the 21 DE genes (**Fig. 3E**).

### Immune landscapes stratify AML patients in independent validation sets and differ across age groups and disease stages

AML is a disease with age-dependent biological specificities^32,33^. Furthermore, pediatric AML are inherently of low immunogenicity and are therefore less likely to respond to single-agent checkpoint inhibition^34^. To characterize the immunological landscape of AML across age groups and longitudinally in patients who initially achieve complete remission (CR) and then experience disease recurrence, we profiled BM samples from a pediatric (CHOP series, n=39 cases) and an adult AML cohort (SAL series, n=38 cases, 58 BM specimens in total). In line with findings in the PMCC discovery series, we identified IFN-dominant, adaptive and myeloid mRNA profiles (**Extended Data Fig. 6A-D**) which individually separated AML cases according to high and low expression values and, when considered in aggregate, dichotomized AML cases into an immune-infiltrated and immune-depleted subtype (**Extended Data Fig. 6E**). As summarized in **Extended Data Fig. 7A**, comparison between children and adults with AML revealed a set of DE immune genes (FDR<4.53×10^−9^) involved with cytokine and chemokine signaling, as indicated by GO (**Extended Data Fig. 8A** and **Supplemental Table 2**) and protein interaction network analysis (**Extended Data Fig. 8B**). Specifically, genes encoding pro-inflammatory and pro-angiogenic chemokines, including *IL8, CCL3L1, CCL3* and *CXCL3*, were expressed at significantly higher levels in adult AML relative to childhood AML (**Extended Data Fig. 7B**). When comparing matched BM samples from adult patients (SAL series) at the time of diagnosis and achievement of CR after induction chemotherapy (**Extended Data Fig. 7C**), we identified a set of DE genes (FDR<5.72×10^−4^) that were enriched for GO biological processes related to the innate immune response, cytokine signaling and toll-like receptor (TLR) cascade (**Supplemental Table 3**). As shown in **Extended Data Fig. 7C**, *FLT3*, Bruton’s tyrosine kinase (*BTK*), IFN regulatory factor 3 (IRF3) and *BMI1*, which have been implicated in leukemia stem cell self-renewal and inhibition of apoptosis, were expressed at significantly lower levels in AML cases in CR, serving as a data reliability check. Finally, immune genes significantly associated with relapsed AML (SAL series) largely captured CD8^+^ T-cell infiltration, elements of T-cell biology, including T-cell receptor (TCR) downstream signaling (*CD3z [CD247], CD3E*), leukocyte differentiation and immune regulation (**Extended Data Fig. 7D** and **Supplemental Table 4**). The increased expression of surrogate markers of terminal T-cell differentiation, senescence and exhaustion [*TBX21* (T-bet)^35,36^, *TIGIT* and *CTLA4*] in relapsed AML suggests that BM-infiltrating cytotoxic T cells may fail to restrain leukemia growth (**Extended Data Fig. 8C**). The DE genes between patient subgroups with newly diagnosed, CR and relapsed AML in the SAL cohort, and between childhood and adult cases, were largely nonoverlapping, as shown in **Extended Data Fig. 7E**.

### IFN-related gene sets improve the prediction of therapy resistance

We then asked whether the IFN-dominant gene module herein computed as the sum of IFN-γ signaling, IFN downstream, immunoproteasome, myeloid inflammation, inflammatory chemokine, IL-10, MAGE, PD-L1 and PD-L2 scores (**Fig. 1** and **Extended Data Fig. 1C**) may assist the prediction of therapeutic resistance, which we empirically defined as failure to achieve CR in patients who survived at least 28 days (primary refractory AML) or as early relapse (<3 months) after achieving CR, as previously published by others^37^. When AML patients in the PMCC cohort were dichotomized based on higher or lower than median IFN scores, a higher percentage of patients with primary refractory disease was observed in the IFN-score^high^ AML cases (65.4% *versus* 34.6%; p=0.0022, Fisher’s exact test), suggesting that transcriptional programs orchestrated by microenvironmental IFN-γ might render AML blasts resistant to chemotherapeutic agents^20,38^. In contrast, the frequency of primary refractory cases was not different when comparing AML patients with higher or lower than median adaptive module scores (29.6% *versus* 25.4%; p=NS) and myeloid module scores (26.7% *versus* 28.1%; p=NS). In multivariate logistic regression analysis, the IFN-related module scores significantly improved the ability of the ELN category to predict therapeutic resistance (**Fig. 4A-B**; AUROC = 0.815 *versus* 0.702 with ELN risk only; model χ^2^=65.87 *versus* 33.43; increased sensitivity compared to ENL risk only=15.6%; increased specificity=3%; decreased false positive rate=11.5%; decreased false negative rate=4.9%), but not patient survival (data not shown). Specifically, the myeloid inflammation score (p=0.003), IFN-γ signaling score (p=0.014) and IFN downstream score (p=0.034) significantly contributed to the model (**Supplemental Table 5**). Gene sets defining gene modules 2 and 3 in **Fig. 1**, reflective of adaptive immune responses and BM infiltration with cells of the myeloid lineage, respectively, were not associated with either therapeutic resistance or patient survival (data not shown).

### *In silico* validation of therapy prediction by immune subtypes in the Beat AML and HOVON cohorts

We next tested the predictive and prognostic power of immune scores *in silico* using a broad collection of public transcriptomic data. We initially devised binary logistic regression models utilizing RNA-sequencing data from 196 patients on the Beat AML Master Trial^®^ with clinical response information^39^. When considering disease type (primary *versus* secondary), WBC count and patient age at diagnosis, the inclusion of genes capturing IFN-γ-related biology significantly improved the predictive ability of the ELN risk category (AUROC=0.921 *versus* 0.709 with ELN cytogenetic risk alone; model χ^2^=106.4 *versus* 29.6; increased specificity=4%; increased sensitivity=17%; decreased false positive rate=39%; decreased false negative rate=18%; **Fig. 4C**).

**Fig. 4:**
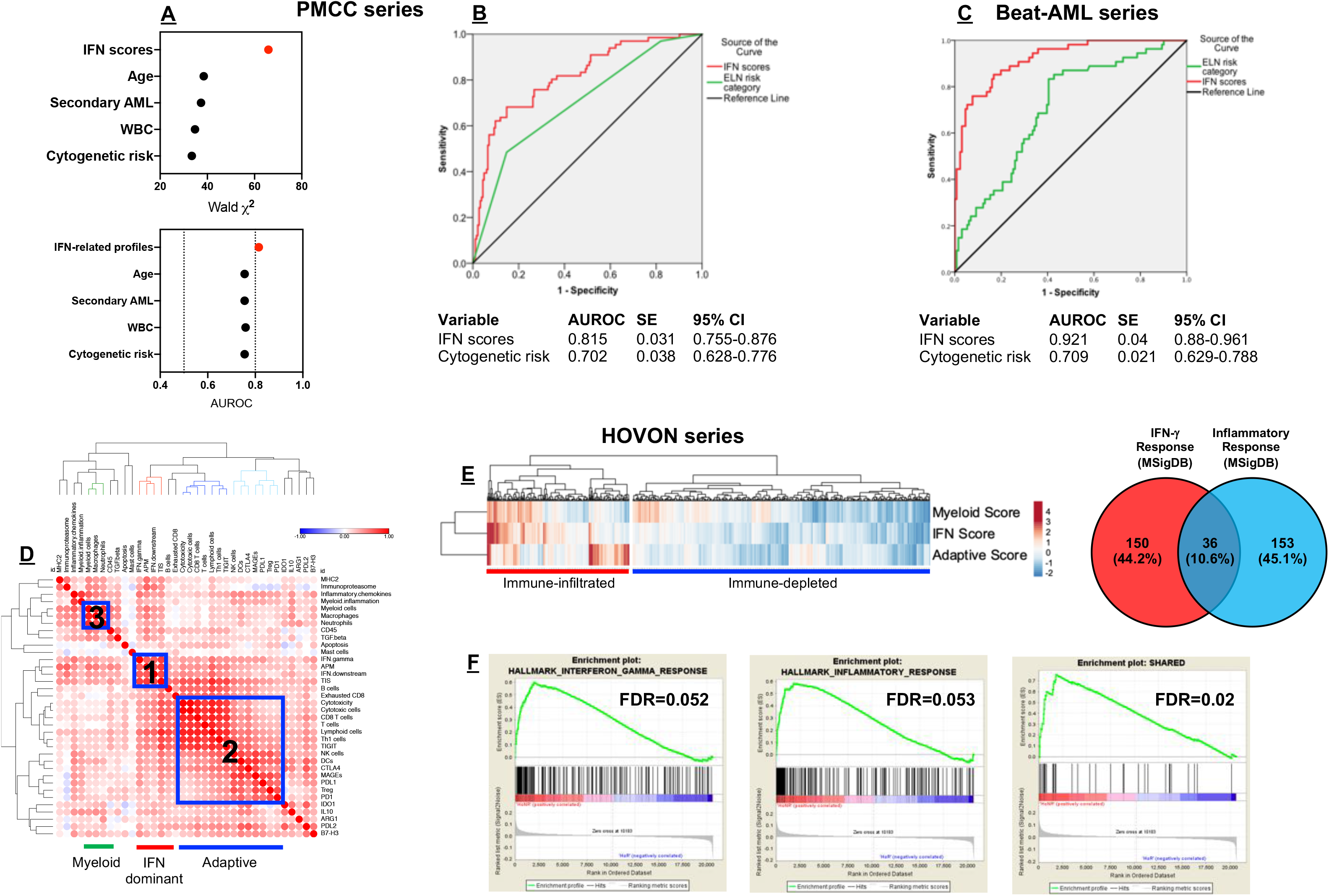
IFN-related mRNA profiles predict therapeutic resistance. **A**) Binary logistic regression predicting therapeutic response from IFN-related scores and conventional prognosticators, i.e., ELN cytogenetic risk category, WBC count at diagnosis, disease type (primary *versus* secondary AML), and patient age at diagnosis (PMCC discovery cohort). AUROC = area under receiver operating characteristic. The dotted line indicates currently accepted thresholds (>0.80) of AUROC with good predictive ability in AML^47^. **B**) AUROC curves measuring the predictive ability of ELN cytogenetic risk and IFN-related scores for therapeutic response (PMCC discovery cohort). SE = standard error; CI = confidence interval. AUROC=1.0 would denote perfect prediction and AUROC=0.5 would denote no predictive ability. **C**) AUROC curves measuring the predictive ability of ELN cytogenetic risk and IFN-related scores for therapeutic response in Beat AML trial specimens (validation cohort). SE = standard error; CI = confidence interval. **D**) Unsupervised hierarchical clustering (Euclidean distance, complete linkage) of the correlation matrix of immune and biological activity signatures identifies co-expression patterns of immune gene sets (correlation value color-coded *per* the legend; Pearson correlation coefficient >0.45; blue boxes) in the bone marrow (BM) microenvironment of AML patients in the HOVON series (n=618 cases with therapy response and ELN cytogenetic risk information). **E**) IFN-dominant, adaptive and myeloid scores in aggregate stratify patients in the HOVON series into immune subtypes (immune-infiltrated and immune-depleted). **F**) Gene set enrichment analysis (GSEA) plots representing the normalized enrichment score (NES) of hallmark IFN-γ-response genes (n=186), inflammatory response genes (n=189) and a subset of overlapping genes (n=36) between IFN-γ and inflammatory gene sets in AML patients in the HOVON series who failed to respond to induction chemotherapy. Gene sets were downloaded from the Molecular Signature Database (MSigDB) and are listed in **Supplemental Table 6**. Each run was performed with 1,000 permutations. FDR=false discovery rate.

Confirming our findings in the PMCC and Beat AML^®^ cohorts, IFN-dominant, adaptive and myeloid mRNA profiles, when used in aggregate, stratified patients in the HOVON database (618 non-promyelocytic AML cases^40^) into subgroups with high and low immune infiltration (**Fig. 4D-E**). Individuals with immune-infiltrated AML had lower leukemia burden (median percentage of BM blasts=56% *versus* 71% in patients with immune-depleted AML; p<0.0001) and tended to have more advanced age at diagnosis (median=51 years, range 15-74, *versus* 46 years, range 15-77; p=0.0067). A higher percentage of patients with IFN-dominant AML failed to achieve CR in response to induction chemotherapy when compared to non-IFN-dominant AML cases (27.2% *versus* 15.2%; p=0.0004, Fisher’s exact test). In contrast, the occurrence of induction failure (IF) was not different when patients were dichotomized based on higher or lower than median adaptive module scores (21.4% *versus* 21.0% IF rate; p=NS) or myeloid module scores (21.7% *versus* 20.7% IF rate; p=NS). Gene set enrichment analysis (GSEA) with all transcripts in the HOVON dataset provided as input and ranked by the log_2_ fold-change between non-responders and responders confirmed the over-expression of curated hallmark gene sets linked to IFN-γ responses and inflammatory responses in chemotherapy-refractory patients (**Fig. 4F; Supplemental Table 6**). When tested in a multinomial logistic regression model incorporating patient age, leukemia burden and ELN cytogenetic risk (available in 615 HOVON cases)^41^, immune gene sets defining the IFN-dominant module significantly and independently predicted whether patients responded to induction chemotherapy and whether they experienced disease relapse (**Supplemental Table 7**). In contrast, immune gene signatures were unable to assist the prediction of nonleukemic deaths (**Supplemental Table 7**).

### Mutations in tumor suppressor genes and transcription factors are enriched in immune-infiltrated AML cases

It has recently been shown that genetic drivers of solid malignancies dictate neutrophil and T-cell recruitment, thus affecting the immune milieu of the tumor and assisting patient stratification^42^. We asked whether clonal driver mutations may correlate with the immune subtypes that we identified herein. We therefore retrieved TCGA AML RNA-sequencing data from cBioPortal (http://www.cbioportal.org/) and computed immune cell type-specific and biological activity scores^22^. The mutational spectrum of TCGA-AML cases, including co-occurrence and exclusivity of the most frequent molecular lesions, is shown in **Extended Data Fig. 9**. Mis-sense mutations, mRNA up-regulation, deep deletion and amplification in IFN downstream genes are summarized in **Extended Data Fig. 10**. Patients with adverse-risk molecular lesions, including somatic *TP53* mutations and *RUNX1* mutations, clustered in the immune-infiltrated subgroup (**Extended Data Fig. 10A**). In particular, IFN-related gene sets, including the tumor inflammation signature (TIS) score, were expressed at significantly higher levels in TCGA-AML cases with *TP53* and *RUNX1* mutations relative to molecular lesions that confer favorable or intermediate risk (**Extended Data Fig. 11A-B**). In contrast, the majority of TCGA-AML cases with *NPM1* mutations with or without *FLT3*-ITD (intermediate-risk and favorable-risk cases, respectively) were classified as immune-depleted. When extending our *in-silico* analysis to the Beat AML^®^ cohort (281 cases in total), 16 out of 17 (94%) *TP53*-mutated AMLs expressed higher levels of genes implicated in downstream IFN signaling and higher levels of *CD8* transcripts and markers of cytotoxicity compared with *TP53* wild-type cases (**Extended Data Fig. 11C-D**).

### IFN-γ-related gene expression profiles correlate with anti-leukemia responses after flotetuzumab immunotherapy

Finally, we hypothesized that higher expression of IFN-γ-related genes in immune-infiltrated AML cases, while underpinning chemotherapy resistance, might identify patients with AML who derive benefit from immunotherapy with flotetuzumab (FLZ; formerly MGD006)^43^, a CD3×CD123 DART^®^. Preliminary anti-leukemic activity of FLZ at a target dose of ≥500ng/kg/day has been recently reported^44,45^. BM samples collected prior to FLZ treatment from 30 adult patients with chemotherapy-refractory or relapsed AML enrolled in the CP-MGD006-01 clinical trial (NCT#02152956) were profiled using the PanCancer IO360™ gene expression assay. Patients’ characteristics are summarized in **Supplemental Table 8**. BM samples from 92% of patients with evidence of FLZ anti-leukemic activity (11 out of 12), which was defined as either CR, CR with partial hematologic recovery (CRh), CR with incomplete hematologic recovery (CRi), partial response or overall benefit (>30% reduction in BM and/or blood blasts), had an immune-infiltrated TME relative to non-responders (**Fig. 5A**). Interestingly, the IFN-dominant module score was significantly higher in patients with chemotherapy-refractory AML compared with relapsed AML at time of FLZ treatment, and in individuals with evidence of anti-leukemic activity compared to non-responders (**Fig. 5B**). Notably, the TIS score was a stronger predictor of anti-leukemic responses to FLZ, with an AUROC value of 0.847 (**Fig. 5B**). On-treatment BM samples (available in 19 patients at the end of cycle 1) displayed increased antigen presentation and immune activation relative to baseline samples, as reflected by higher TIS scores (6.47±0.22 *versus* 5.93±0.15, p=0.0006), antigen processing machinery scores (5.67±0.16 *versus* 5.31 ±0.12, p=0.002), IFN-γ signaling scores (3.58±0.27 *versus* 2.81±0.24, p=0.0004) and *PD-L1* expression (3.43±0.28 *versus* 2.73±0.21, p=0.0062; **Fig. 5C**). Overall, these data substantiate a clinical benefit for AML patients with an immune-infiltrated TME, support a local immune-modulatory effect of FLZ treatment and validate the translational relevance of our findings.

**Fig. 5:**
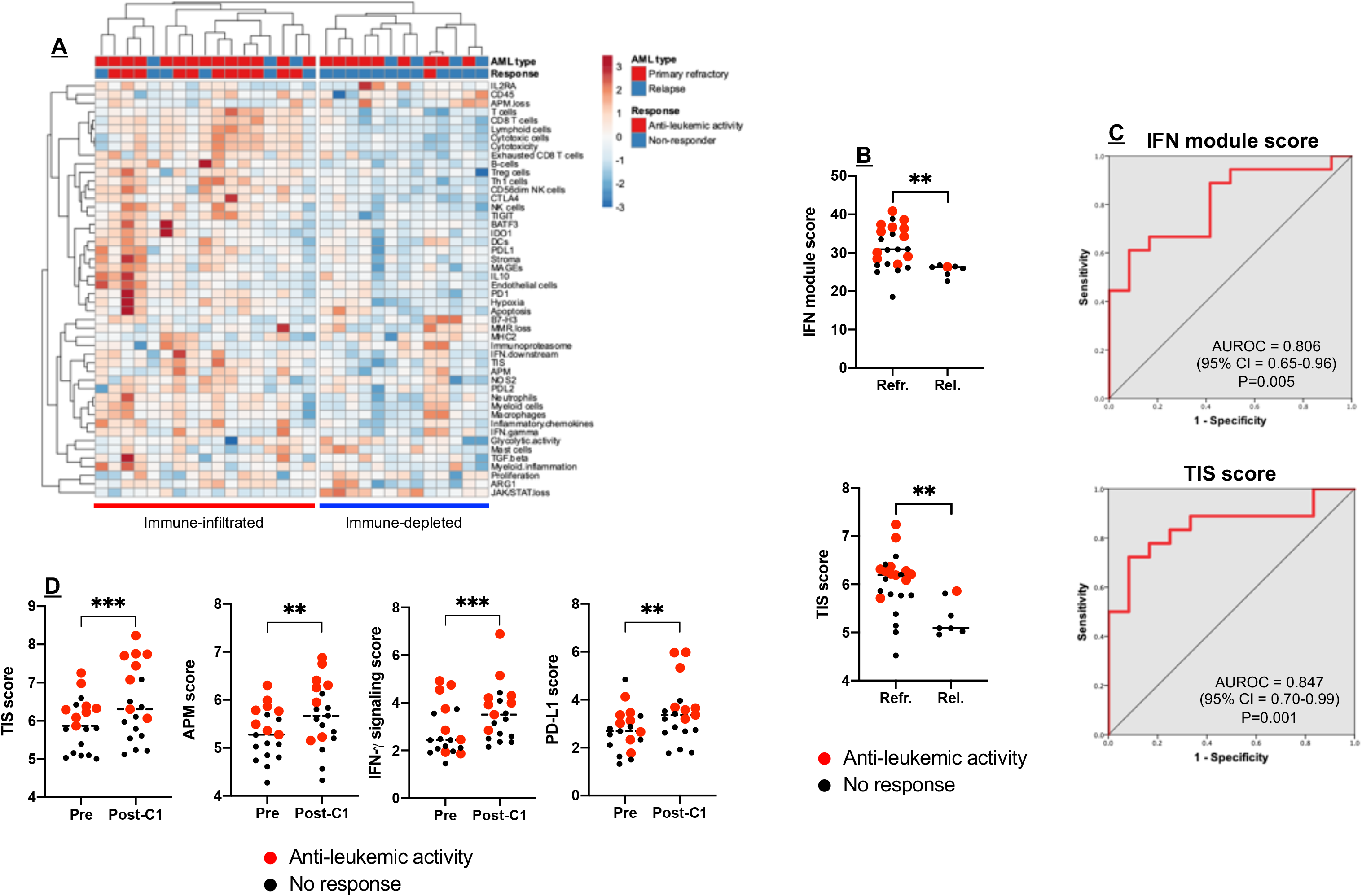
Immune subtypes associate with response to flotetuzumab immunotherapy. **A**) Unsupervised hierarchical clustering (Euclidean distance, complete linkage) of immune and biological activity signatures in the bone marrow (BM) microenvironment of patients with relapsed/refractory AML (n=30) receiving flotetuzumab (FLZ) immunotherapy in the CP-MGD006-01 clinical trial (NCT#02152956). Anti-leukemic response was defined as either complete remission (CR), CR with incomplete hematologic recovery (CRi), CR with partial hematologic recovery (CRh), partial remission (PR) or “other benefit” (OB; >30% decrease in BM blasts). Non-responders were individuals with either treatment failure (TF), stable disease (SD) or progressive disease (PD). Chemotherapy refractoriness was defined as ≥2 induction attempts or 1^st^ CR with initial CR duration <6 months. **B**) IFN module score and TIS score in baseline BM samples from patients with primary refractory and relapsed AML. Red dots denote patients with evidence of FLZ anti-leukemic activity. Horizontal lines indicate median values. Comparisons were performed with the Mann-Whitney *U* test for unpaired data (two-sided). **P<0.01. **C**) Area under receiver operating characteristic (AUROC) curves measuring the predictive ability of the IFN-module score and tumor inflammation signature (TIS) for therapeutic response to FLZ. CI=confidence interval. **D**) Immune activation in the TME during FLZ treatment (matched BM samples from 19 patients). Red dots denote patients with evidence of FLZ anti-leukemic activity. Horizontal lines indicate median values. Comparisons were performed with the Mann-Whitney *U* test for paired data (two-sided). Pre = baseline. C1 = cycle 1. **P<0.01. ***P<0.001.

## Discussion

Using large cohorts of subjects, the current study is the first to reveal underlying transcriptomic features that stratify the TME of AML into immune subtypes and may assist therapeutic predictions by defining patients who will potentially derive the greatest benefit from immunotherapies^11,46^. We identified two subtypes of differentially-infiltrated tumors, an observation that was validated in independent childhood and adult AML series, reinforcing the notion that unique molecular features can distinguish AML across age groups^33^. In agreement with commonly accepted criteria that AUROCs of 0.8-0.9 indicate good predictive ability, the IFN-related gene sets identified in our study improved the prediction of therapeutic resistance following conventional ‘3+7’ cytarabine and anthracycline chemotherapy beyond that provided by the ELN cytogenetic risk category (AUC=0.815 in PMCC cases [discovery series] and 0.870 in Beat AML^®^ cases [*in silico* validation series])^47^. In recent Southwestern Oncology Group (SWOG) and MD Anderson Cancer Center clinical trials, pretreatment covariates such as cytogenetic risk and age only yielded AUROCs of 0.65 and 0.59 for therapeutic resistance, respectively^37^. Our models incorporating IFN-related mRNA profiles also outperform a recently developed 29-gene and cytogenetic risk predictor of chemotherapy resistance (AUROC=0.76)^48^. Intriguingly, an IFN-related DNA damage resistance signature (IRDS) correlates with resistance to adjuvant chemotherapy and with recurrence after radiotherapy in patients with breast cancer^38^, suggesting that tumor cells over-expressing IRDS genes, including *STAT1, ISG15* and *IFIT1*, as a result of chronic activation of the IFN signaling pathway might receive pro-survival rather than cytotoxic signals in response to DNA damage^49^.

The immune-infiltrated AML cases in the PMCC cohort were highly immune-suppressed, as indicated by elevated expression of IFN-inducible negative immune checkpoints and immunotherapy targets *IDO1* and *PD-L1*. Furthermore, adults with relapsed AML in the SAL series expressed higher levels of T-cell exhaustion molecules relative to matched pretreatment samples, suggesting the occurrence of escape from immune surveillance at the time of disease relapse^50^. In general, solid tumors with a substantial T-cell component and displaying a type I immune response are associated with better OS and progression-free survival estimates^6^. However, the highly proliferative IFN-γ-dominant solid tumors may correlate with a less favorable survival, despite being infiltrated with CD8^+^ T cells and harboring the highest type 1 macrophage signature scores^6^. By utilizing targeted IGEP, our study highlights that the activation of IFN-related pathways and the relative abundance of immune cell types, including the over-expression of T-cell markers and TCR signaling intermediates in relapsed AML relative to disease onset, have negative prognostic implications in AML; this is conceivably the result of a non-productive anti-leukemia immune response and/or IFN-driven resistance to DNA damage induced by chemotherapeutic agents^20^.

The heterogeneity of immune infiltration can also be determined by tumor cell-intrinsic factors, including chemokine secretion^51^ and expression of cancer driver genes, all of which affect response to immunotherapies^42,52,53^. Interestingly, we detected associations between mutations in tumor suppressor and cancer driver genes and immune subtypes of AML and, for the first time, we identified *TP53* mutations as being strongly correlated with an IFN-γ dominant TME and with prognostic protein signatures, including the expression of *PD-L1, FOXP3*, cytotoxicity markers and *PTEN*, that were revealed by spatially-resolved, highly multiplexed protein profiling. This observation is backed by a recent study showing higher proportions of PD-L1-expressing CD8^+^ T cells, activation of IFN-γ-associated genes and favorable responses to pembrolizumab immunotherapy in *TP53*-mutated lung cancers^54^. It is tempting to speculate that immune-infiltrated, *TP53*-mutated AML cases, which have very low response rates when treated with standard anthracycline-based and cytarabine-based induction chemotherapy, could benefit from T cell-targeting approaches and/or hypomethylating agents that potentially alter the immune surveillance of AML.^55^ Finally, IFN-γ-related gene expression programs in the AML TME, including the TIS score, predicted response to immunotherapy with FLZ in 30 heavily pre-treated patients with relapsed/refractory AML on clinical trial CP-MGD006-01. FLZ treatment was associated with increased expression of antigen processing machinery genes, enhanced IFN-γ signaling and heightened *PD-L1* expression. The latter finding provides a biological rationale for designing clinical studies with sequential FLZ and checkpoint inhibitor blockade in AML patients in remission with minimal residual disease. In conclusion, our work unveils the heterogeneity of the immune landscape of AML and provides a novel precision medicine-based conceptual framework for delivering T cell-targeting immunotherapy to subgroups of patients with IFN-γ-dominant AML, who may be refractory to conventional cytotoxic chemotherapy but responsive to T-cell engagers. The immunological stratification of pre-treatment BM samples may therefore enable rapid risk prediction and selection of frontline therapeutic modalities^11^, in conjunction with cytogenetic and mutational information.

## Methods

Methods, including statements of data availability and any associated accession codes and references, are available online.

## Disclosure of potential conflict of interest

The authors have no potential conflicts of interest to disclose.

## Authors’ Contributions

**Concept and design**: S. Rutella

**Development of methodology**: J. Vadakekolathu, T. Hood, S.E. Church, S. Reeder, S.E. Warren, Y. Liang

**Acquired, consented and managed patients; processed patient samples**: M.D. Minden, N. Ibrahimova, A. Arruda, J. Muth, P.J.M. Valk, B. Löwenberg, M. Bornhäuser, S.K. Tasian, M.P. Rettig, J.F. DiPersio

**Analysis and interpretation of data**: J. Vadakekolathu, M.D. Minden, T. Hood, S.E. Church, S. Reeder, A.H. Sullivan, E. Viboch, S.E. Warren, Y. Liang, M. Schmitz, A. Cesano, P.J.M. Valk, B. Löwenberg, A.G. Pockley, M. Bornhäuser, S.K. Tasian, J. Davidson-Moncada, J.F. DiPersio, S. Rutella

**Clinical trial implementation**: J.F. DiPersio was principal investigator at Washington University in St. Louis, St. Louis, United States of America. B. Löwenberg was principal investigator at Erasmus University Medical Centre, Rotterdam, Netherlands.

**Writing of the manuscript**: S. Rutella

**Review and/or revision of the manuscript**: J. Vadakekolathu, M.D. Minden, T. Hood, S.E. Church, E. Viboch, S.E. Warren, M. Schmitz, A. Cesano, P.J.M. Valk, B. Löwenberg, A.G. Pockley, M. Bornhäuser, S.K. Tasian, M.P. Rettig, J. Davidson-Moncada, J.F. DiPersio, S. Rutella

**Study supervision**: S. Rutella

## Acknowledgements

This work was supported by grants from the Qatar National Research Fund (NPRP8-2297-3-494 to S. Rutella), the Roger Counter Foundation, United Kingdom (to A.G. Pockley and S. Rutella), the John and Lucille van Geest Foundation (to A.G. Pockley and S. Rutella), the James Skillington Challenge for Leukemia (to A.G. Pockley), the NCI K08 CA184418 and the Andrew McDonough B+ Foundation (to S.K. Tasian). The Study Alliance of Leukemia (www.sal-aml.org) is gratefully acknowledged for providing primary patient material and clinical data.

## Online Methods

### Patients’ demographics (discovery cohorts)

Patient and disease characteristics are detailed in Table 1. Primary patient specimens (non-promyelocytic AML) and associated clinical data were obtained on research protocols approved by the Investigational Review Boards of the Children’s Hospital of Philadelphia (CHOP), USA and Princess Margaret Cancer Centre (PMCC), Canada and by the Ethics Committee of TU Dresden and Studienallianz Leukämie (SAL), Germany.

### Patients’ demographics (immunotherapy cohort)

Thirty patients with primary refractory (n=23) and relapsed AML (n=7) treated with flotetuzumab (FLZ) at the recommended phase 2 dose (500 ng/kg/day) on the CP-MGD006-01 clinical trial (NCT#02152956) were included in this study. Patient and disease characteristics are detailed in **Supplemental Table 8**. BM aspirates were collected at baseline (n=30) and after cycle 1 of FLZ (n=19) to evaluate the temporal immunological effects associated with therapeutic response. Patients received a lead-in dose of FLZ during week (W) 1, followed by 500 ng/kg/day during weeks 2-4 of cycle 1, and a 4-day on/3-day off schedule for cycle 2 and beyond. Disease status was assessed by modified IWG criteria.

### RNA isolation and processing

Messenger RNA was isolated and processed as previously described^41^. For the PMCC, SAL and CHOP patient cohorts, approximately 100 ng per sample of RNA extracted from 387 bulk BM aspirates from AML patients treated with curative intent were analyzed on the nCounter™ FLEX analysis system using the PanCancer Immune [PCI] profiling panel (for research use only and not for use in diagnostic procedures), which measures mRNA levels of 740 genes representing 24 immune cell types, common checkpoint inhibitors, cancer testis antigens and genes covering both the innate and adaptive immune response (**Extended Data Fig. 2B**)^25,56,57^. BM samples from patients receiving FLZ immunotherapy were analyzed using the PanCancer IO360™ mRNA panel (for research use only and not for use in diagnostic procedures).

### nCounter data quality control, data normalization and signature calculation

The reporter probe counts, i.e., the number of times the color-coded barcode for that gene is detected, were tabulated in a comma separated value (CSV) format for data analysis with the nSolver™ software package (version 4.0.62) and nSolver Advanced Analysis module (version 2.0.115; NanoString Technologies). The captured transcript counts were normalized to the geometric mean of the housekeeping reference genes included in the assay and the code set’s internal positive controls.

The relative abundance of immune cell types and immuno-oncology biological signatures were computed as previously published^22,23^. For samples run on the PCI profiling panel, we also calculated an approximation of the Tumor Inflammation Signature (TIS) using 16 of the 18 functional genes and 5 of the 10 housekeeper genes that are present in the PCI profiling panel^18^.

### Gene ontology (GO) and gene set enrichment analysis (GSEA)

Metascape.org was used to enrich genes for GO biological processes and pathways. GSEA was performed using the GSEA software v.3.0 (Broad Institute, Cambridge, USA)^58^. Hallmark IFN-γ response and inflammatory response gene sets (M5913) were downloaded from the Molecular Signature Database (MSigDB)^22,23^.

### GeoMx™ Digital Spatial Profiling (DSP)

Ten FFPE BM biopsies from patients with newly diagnosed AML (SAL series) were profiled using the GeoMx DSP platform. Samples were stained using 3 fluorescent visualization markers, CD3 (T cell), CD123 (myeloid blast), SYTO™ 83 (nuclei), and 31 UV-cleavable oligo-labeled antibodies (**Supplemental Table 9**). Stained slides were loaded on the DSP instrument and digitally scanned. Fluorescent scans were used to select 24 geometric regions of interest (ROIs)^27,59^. The DSP instrument then UV-illuminated selected ROIs to release conjugated oligos and the micro-capillary fluidics system collected released oligos, which were counted on the nCounter system.

### Data sources for *in silico* analyses

The first data series (E-MTAB-3444), hereafter referred to as the HOVON series^40^, was retrieved from Array Express and encompassed three independent cohorts of adults (≤60 years) with *de novo* AML. BM and blood samples were collected at diagnosis and were analyzed on the Affymetrix Human Genome U133 Plus 2.0 Microarray^40,60^. Patients were treated with curative intent according to the Dutch-Belgian Hematology-Oncology Cooperative Group and the Swiss Group for Clinical Cancer Research (HOVON/SAKK) AML-04, −04A, −29, −32, −42, −42A, −43 or −92 protocols (available at http://www.hovon.nl). Clinical annotations were provided by the authors. The second data series, hereafter referred to as The Cancer Genome Atlas (TCGA) series, consisted of RNA-sequencing data (Illumina HiSeq2000) from 162 adult AML patients with complete cytogenetic, immunophenotypic and clinical annotation who were enrolled on Cancer and Leukemia Group B (CALGB) treatment protocols 8525, 8923, 9621, 9720, 10201 and 19808^61^. RNA and clinical data were retrieved from the TCGA data portal (https://tcga-data.nci.nih.gov/tcga/tcgaDownload.jsp). The third data series (Beat AML) was retrieved using the VIZOME user interface (http://www.vizome.org/aml/) and consisted of RNA-sequencing data from primary specimens from 242 AML patients with detailed clinical annotations, including diagnostic information, treatments, responses and outcomes treated on the Beat AML Master Trial^39^. Patient and disease characteristics for *in silico* data sources are summarized in **Supplemental Table 10**.

### Statistical analyses

Descriptive statistics included calculation of mean, median, SD, and proportions to summarize study outcomes. Comparisons were performed with the Mann-Whitney U test for paired or unpaired data (two-sided), as appropriate, or with the ANOVA with correction for multiple comparisons. IBM SPSS Statistics (version 24) and GraphPad Prism (version 8) were used for statistical analyses. A two-tailed *p* value <0.05 was considered to reflect statistically significant differences. The log-rank (Mantel-Cox) test was used to compare survival distributions.

Therapeutic resistance was defined as failure to achieve complete remission (CR) despite not experiencing early treatment-related mortality (within 28 days of chemotherapy initiation; primary refractory cases) or as early relapse (<3 months) after achieving CR^37^. Overall survival (OS) was computed from the date of diagnosis to the date of death. Relapse-Free-Survival (RFS) was measured from the date of first CR to the date of relapse or death. Subjects lost to follow-up were censored at their date of last known contact.

Binary logistic regression and multinomial logistic regression were used to ascertain the relative contribution of immune subtypes and other pretreatment covariates selected *a priori* based on known clinical relevance (ELN risk group, *FLT3*-ITD status, *NPM1* mutational status, patient age at diagnosis and primary *versus* secondary AML) toward the predicted likelihood of response to induction chemotherapy, AML relapse and patient death^41^. Variables measured after the initiation of induction chemotherapy were excluded, since the goal of the current study was to identify AML patients at higher risk of treatment failure prior to starting therapy.

### Data availability

Gene expression data have been deposited in NCBI’s Gene Expression Omnibus^62^ and are accessible through GEO Series accession number GSE134589 (https://www.ncbi.nlm.nih.gov/geo/query/acc.cgi?acc=GSE134589). Processed input data and basic association analyses will be made available from the corresponding author on request for the purpose of conducting legitimate scientific research.

---

**Extended Data Fig. 1.**
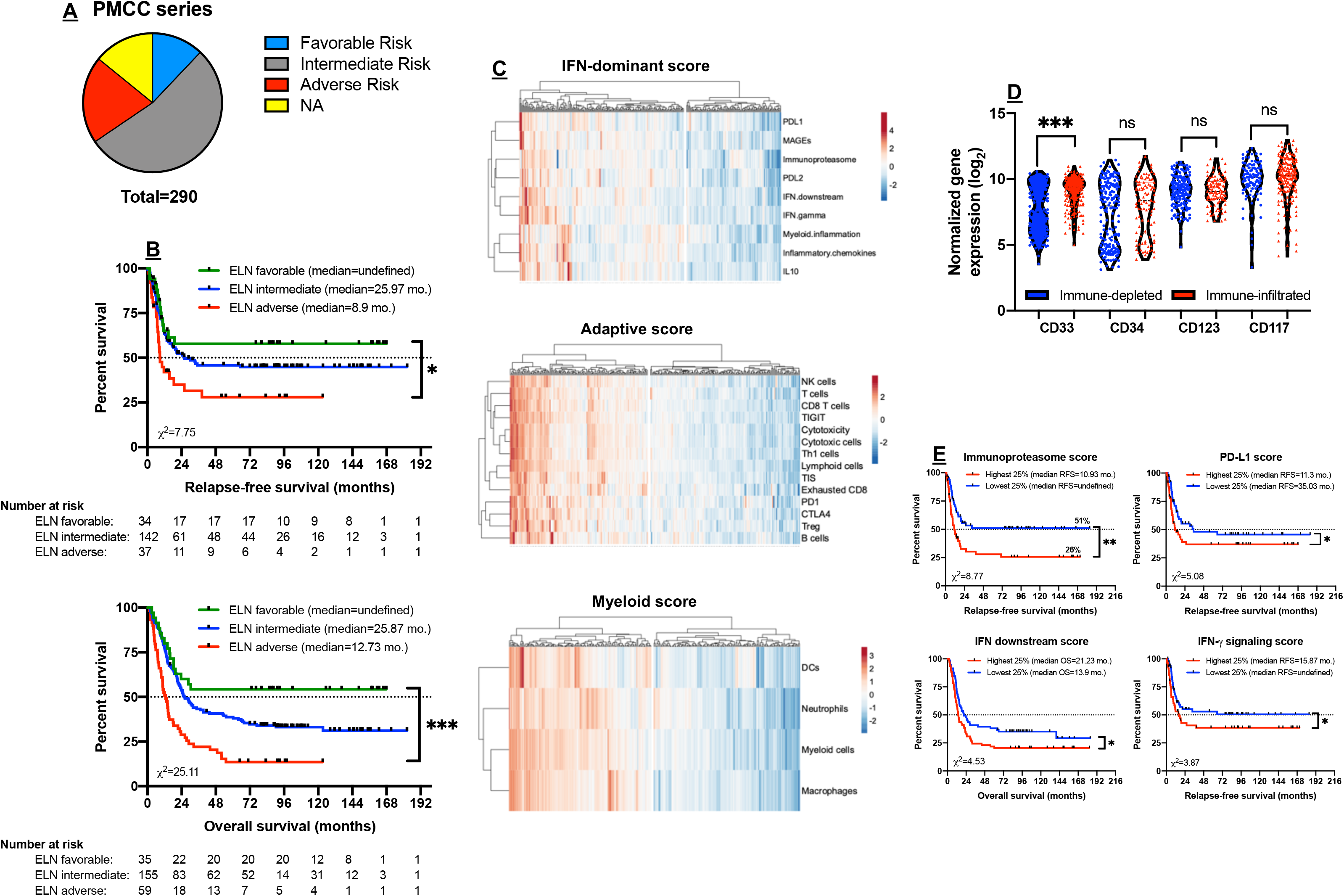

**Extended Data Fig. 2.**
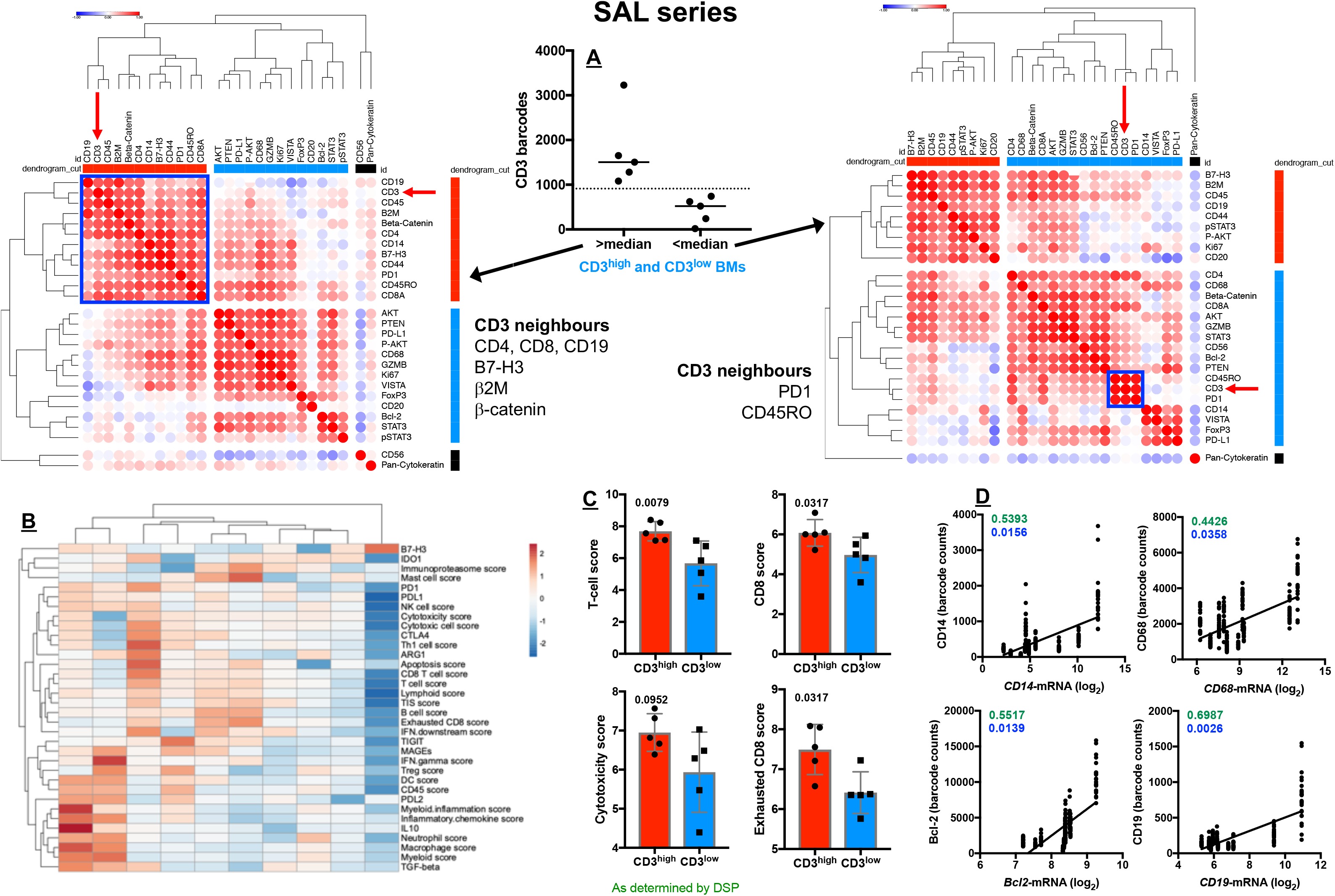

**Extended Data Fig. 3.**
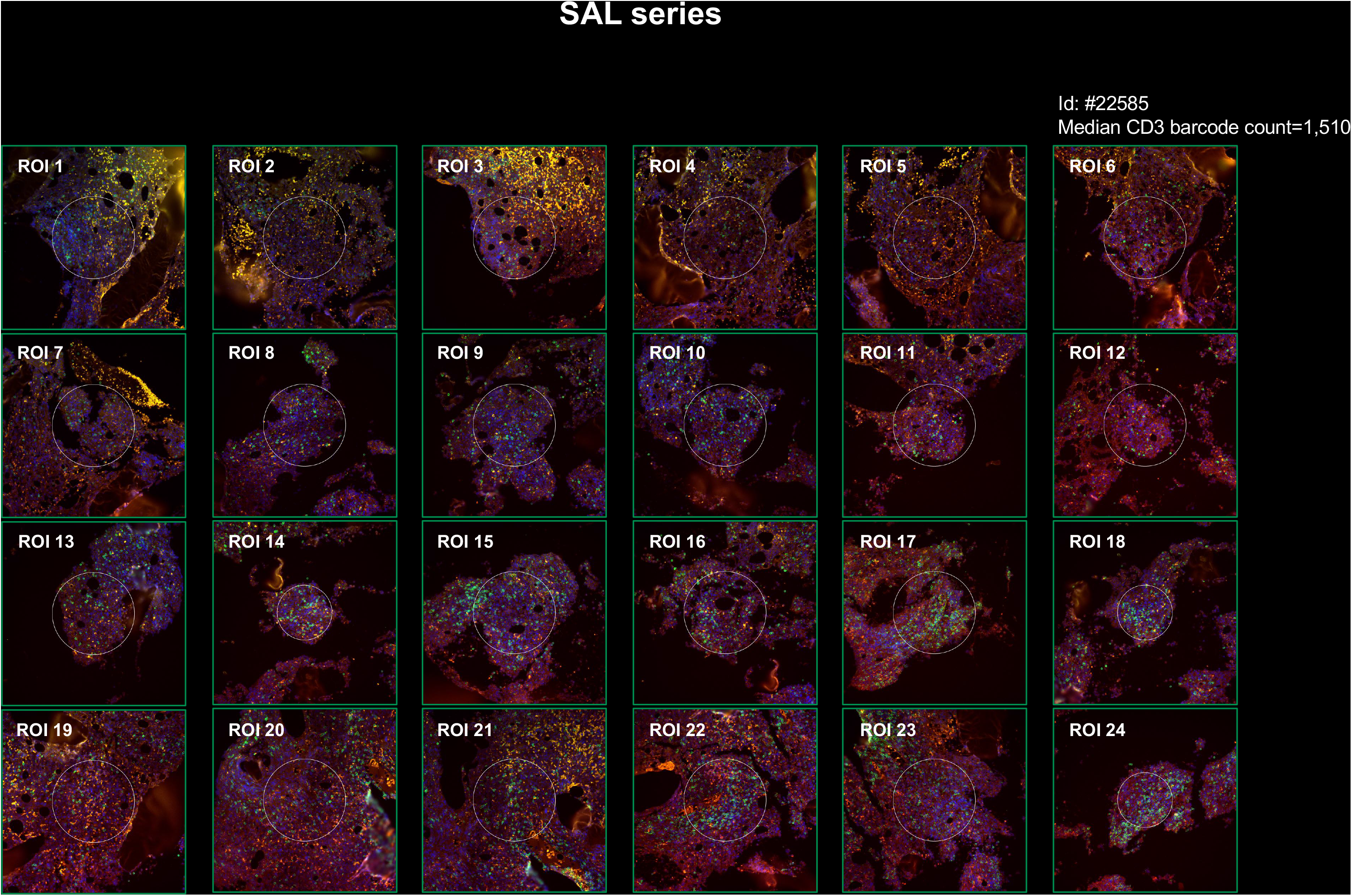

**Extended Data Fig. 4.**
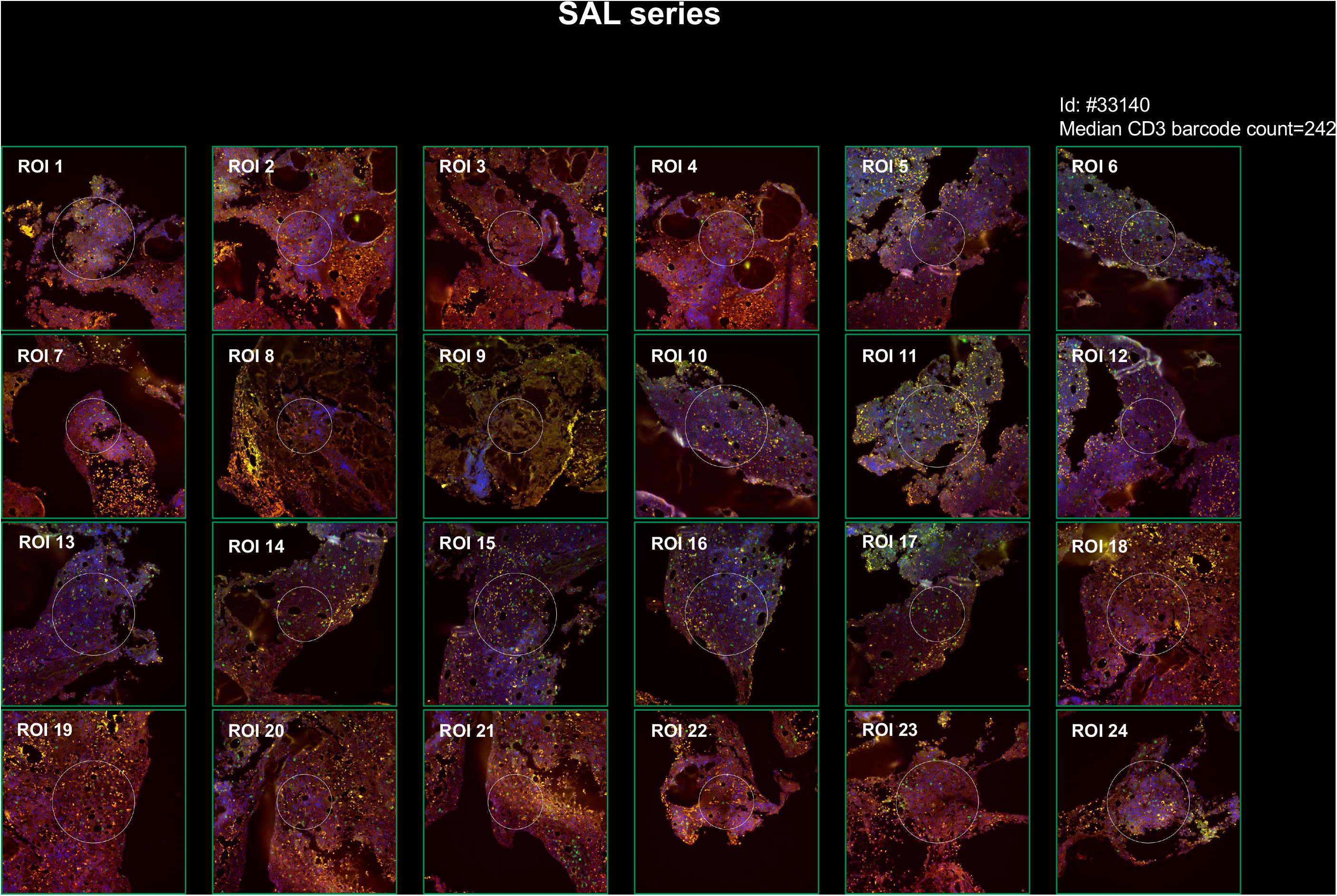

**Extended Data Fig. 5.**
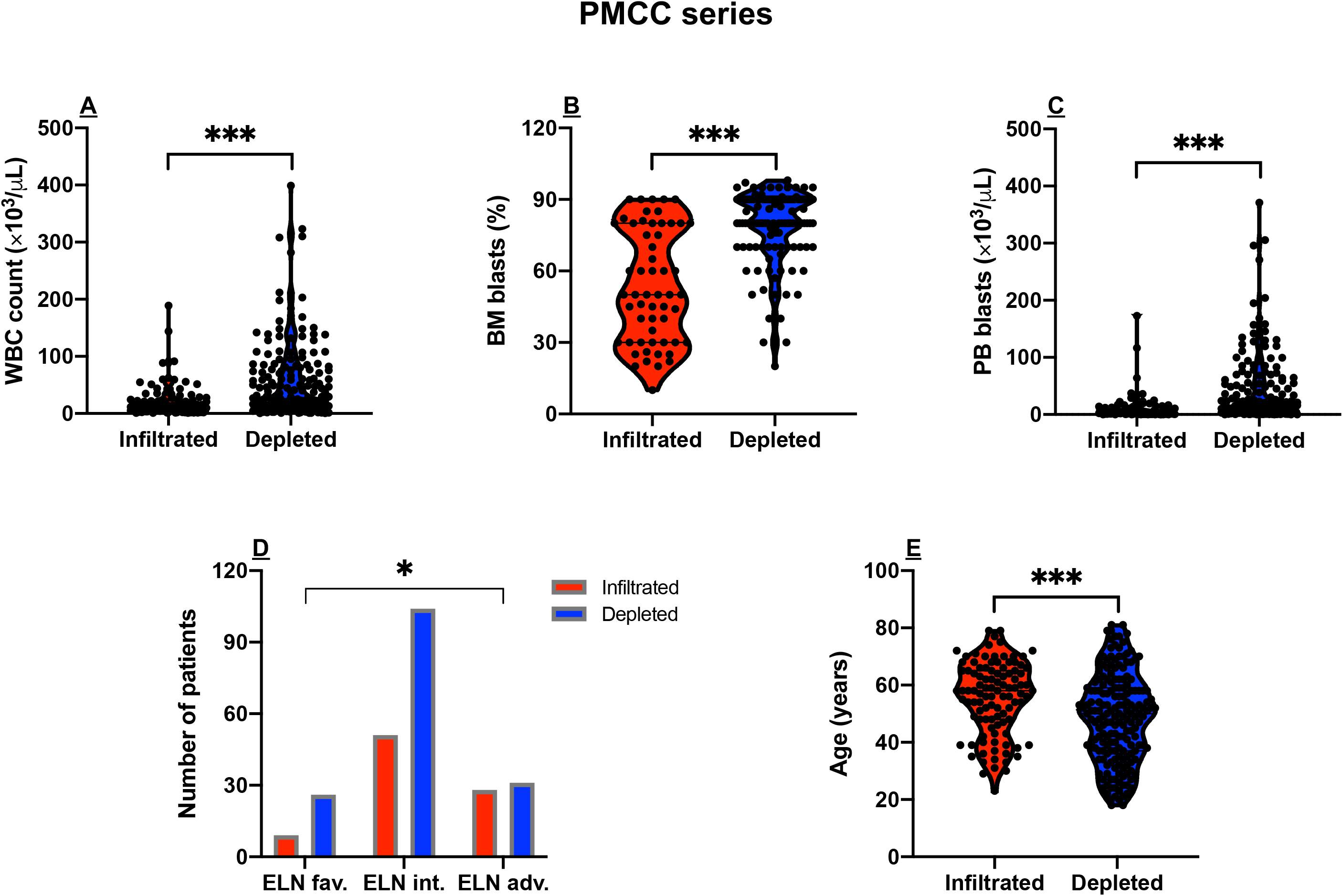

**Extended Data Fig. 6.**
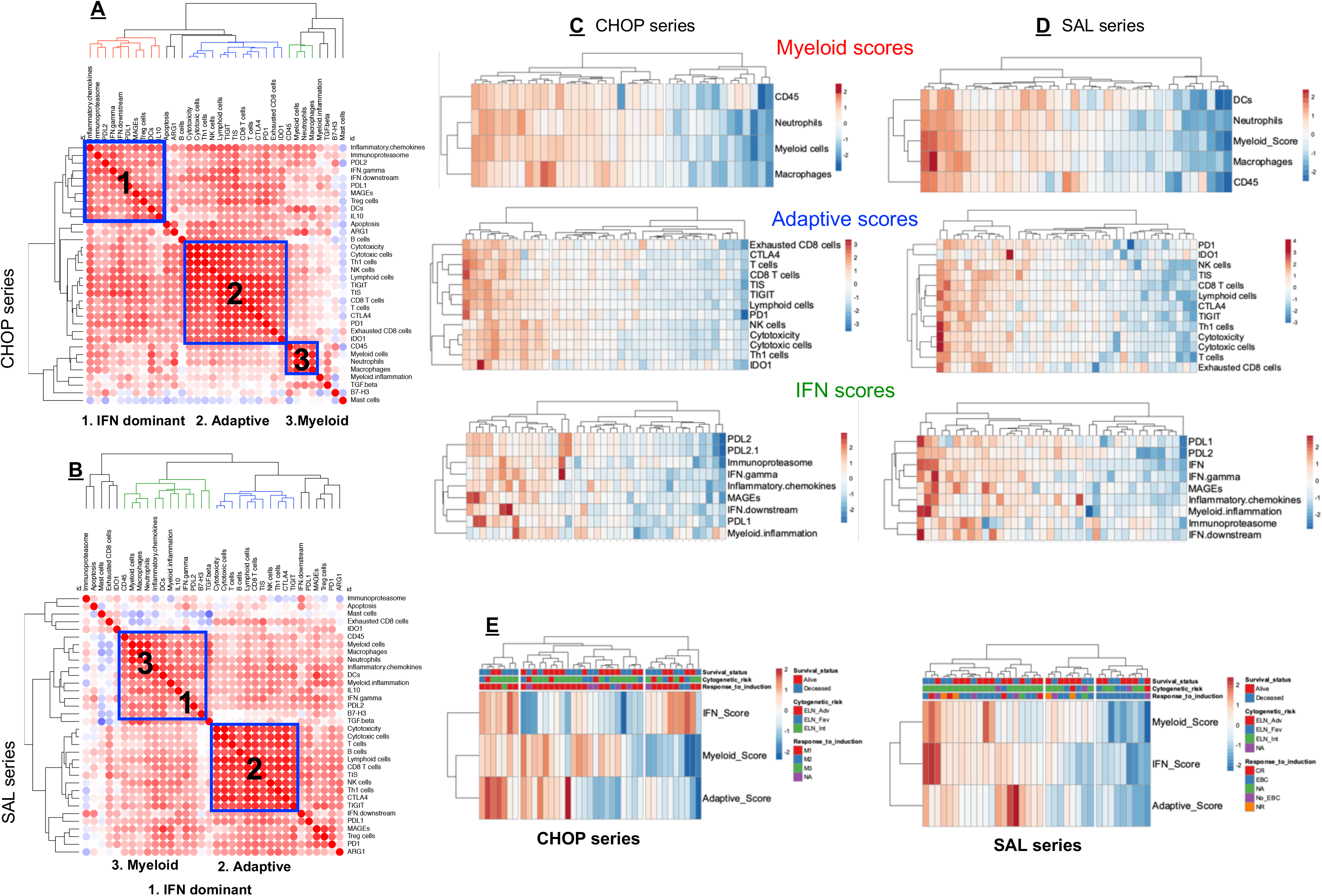

**Extended Data Fig. 7.**
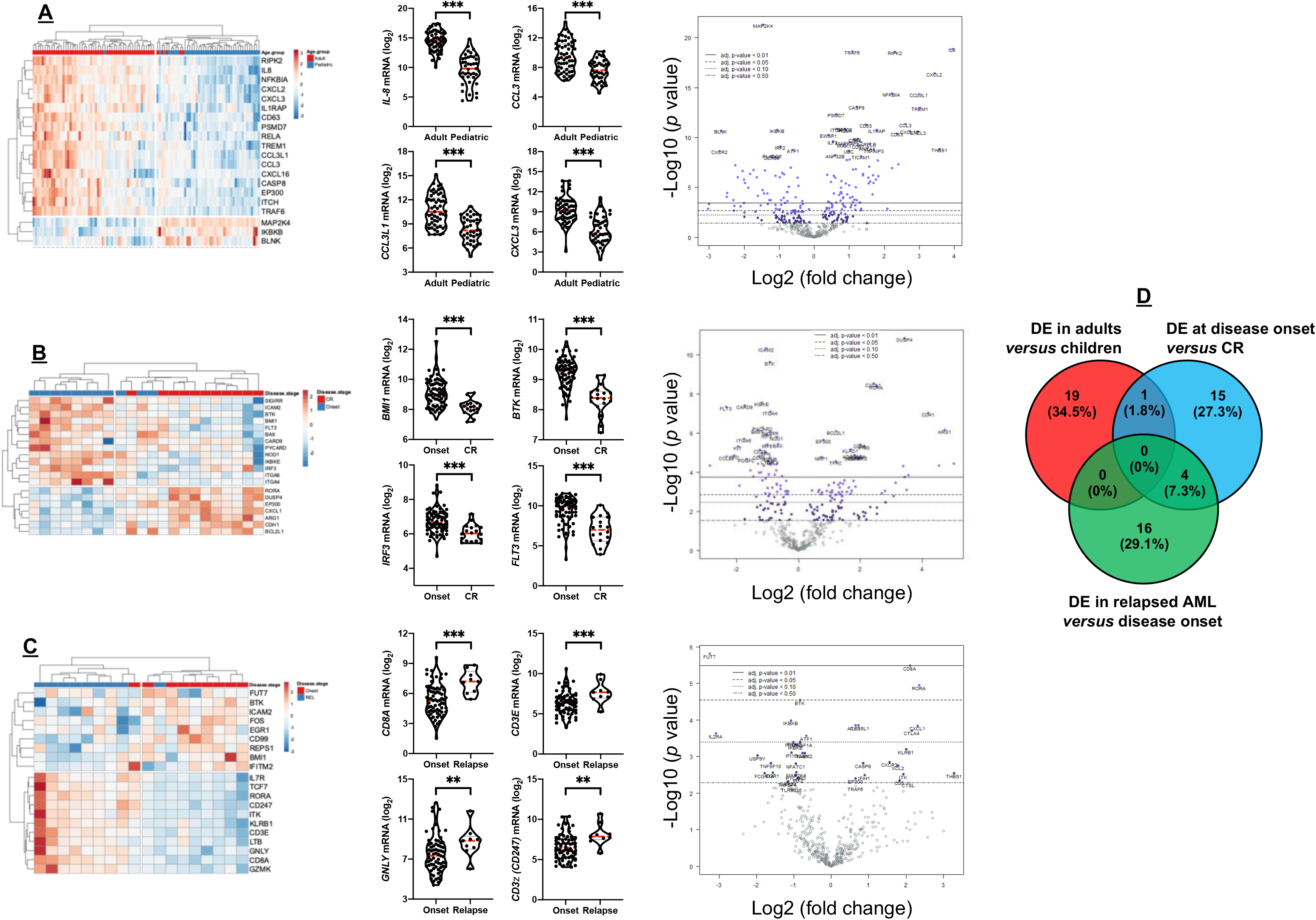

**Extended Data Fig. 8.**
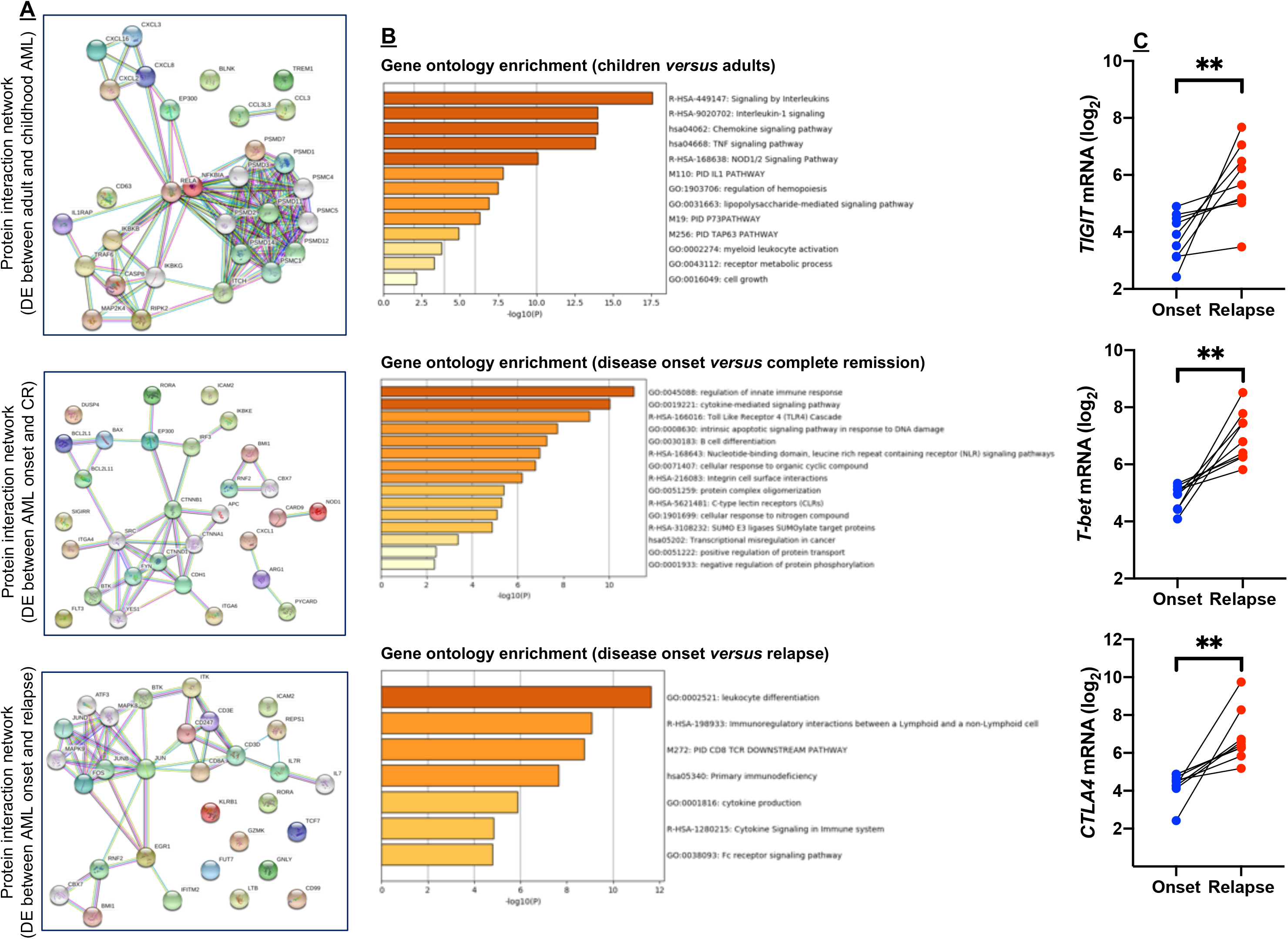

**Extended Data Fig. 9.**
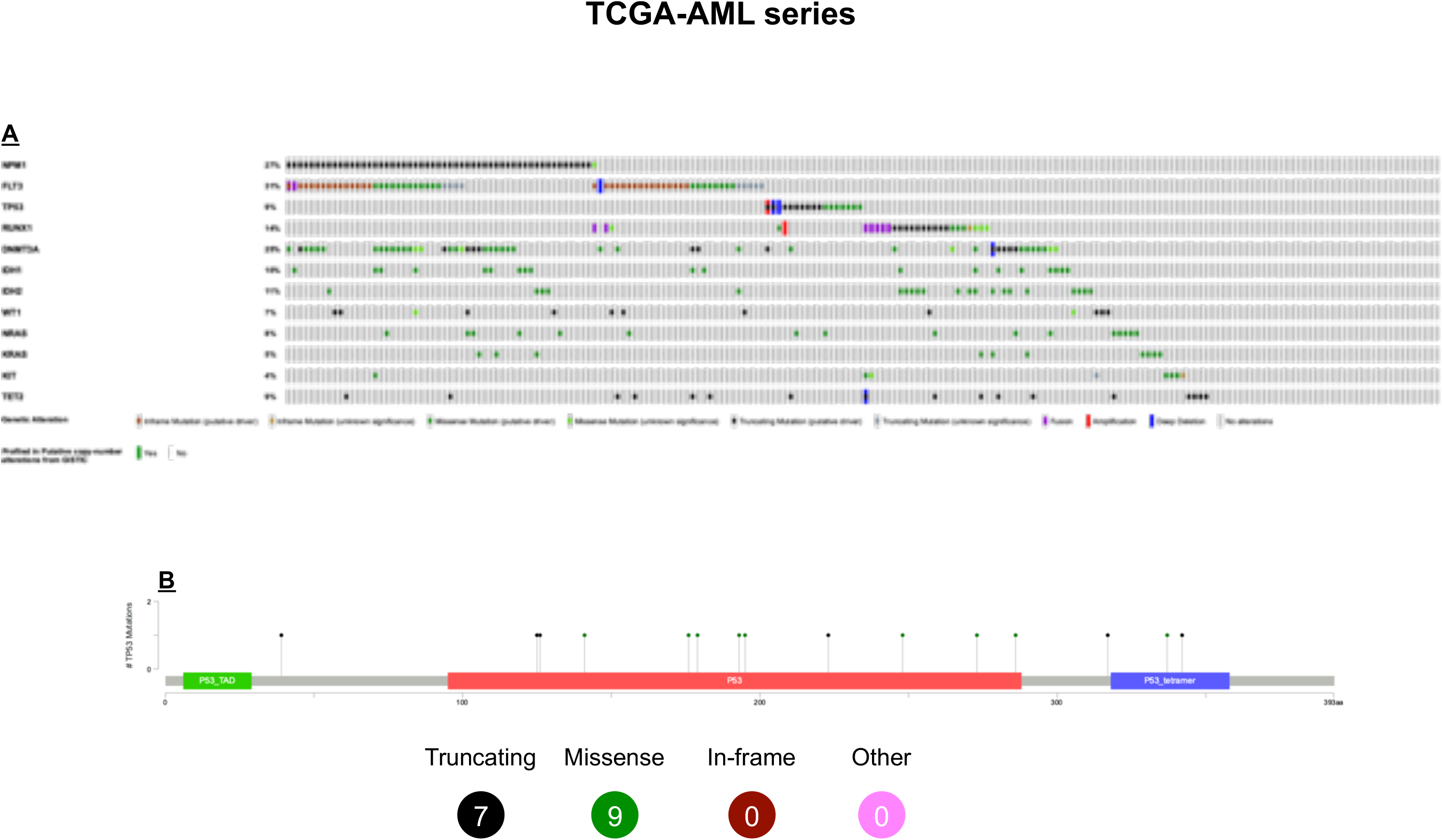

**Extended Data Fig. 10.**
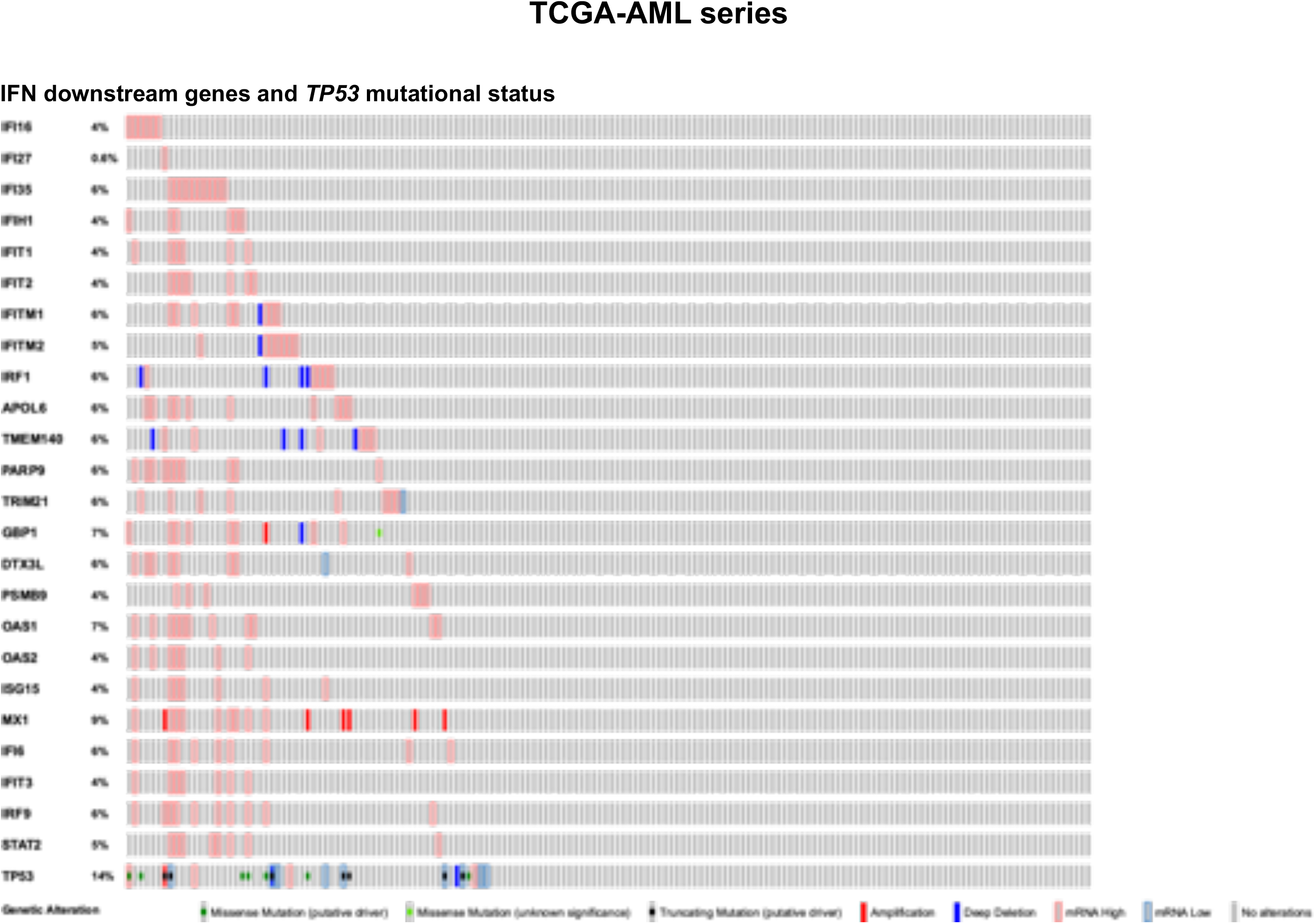

**Extended Data Fig. 11.**
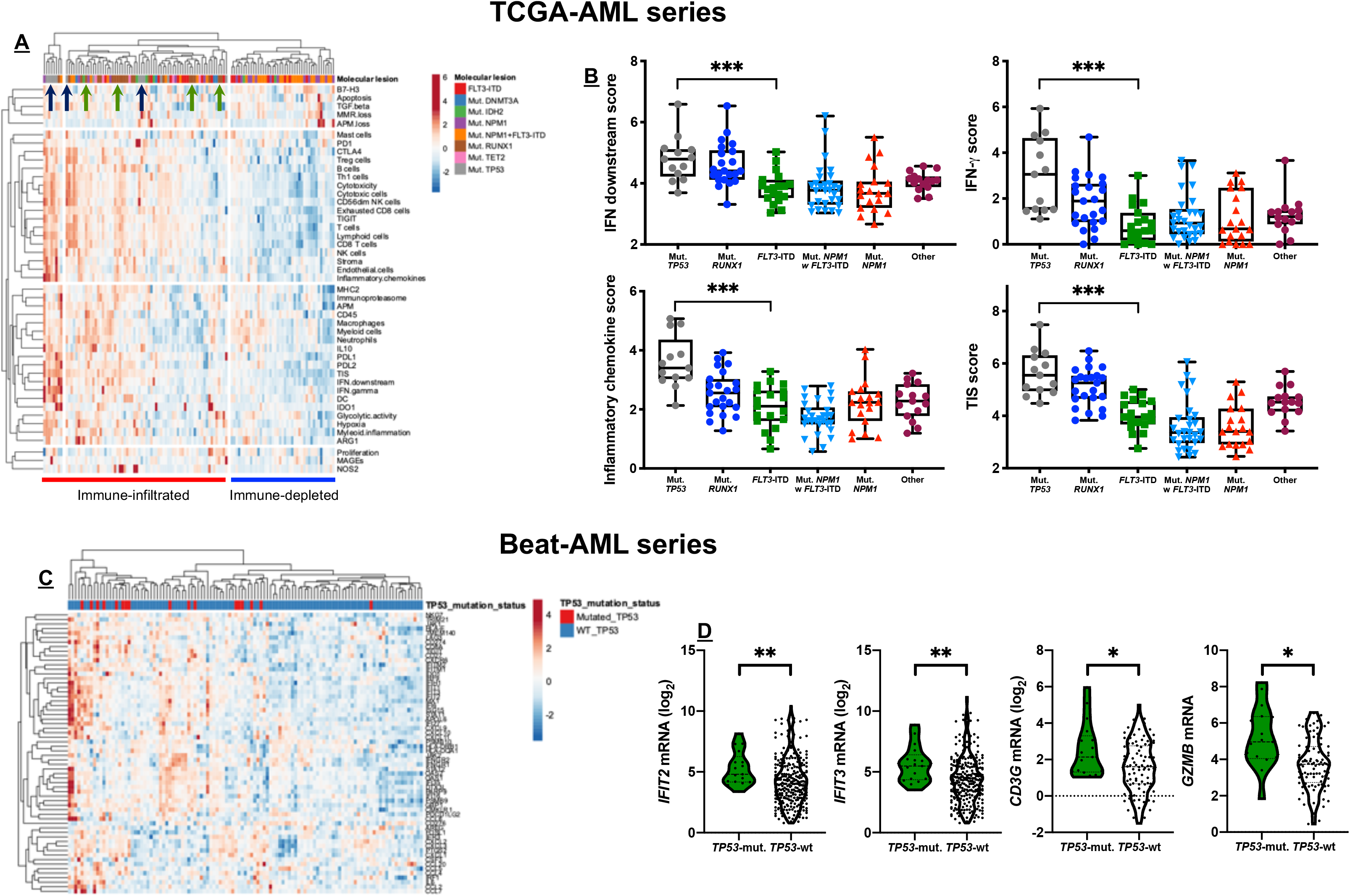

## References

1. Dohner, H., Weisdorf, D.J. & Bloomfield, C.D. Acute myeloid leukemia. N Engl J Med 373, 1136–1152 (2015).

2. Papaemmanuil, E., et al. Genomic classification and prognosis in acute myeloid leukemia. N Engl J Med 374, 2209–2221 (2016).

3. Khwaja, A., et al. Acute myeloid leukaemia. Nat Rev Dis Primers 2, 16010 (2016).

4. Coombs, C.C., Tallman, M.S. & Levine, R.L. Molecular therapy for acute myeloid leukaemia. Nat Rev Clin Oncol 13, 305–318 (2016).

5. Binnewies, M., et al. Understanding the tumor immune microenvironment (TIME) for effective therapy. Nat Med 24, 541–550 (2018).

6. Thorsson, V., et al. The immune landscape of cancer. Immunity 48, 812–830 (2018).

7. Gill, S., et al. Preclinical targeting of human acute myeloid leukemia and myeloablation using chimeric antigen receptor-modified T cells. Blood 123, 2343–2354 (2014).

8. Gibney, G.T., Weiner, L.M. & Atkins, M.B. Predictive biomarkers for checkpoint inhibitor-based immunotherapy. Lancet Oncol 17, e542–e551 (2016).

9. Cristescu, R., et al. Pan-tumor genomic biomarkers for PD-1 checkpoint blockade-based immunotherapy. Science 362 (2018).

10. Daver, N., et al. Efficacy, safety, and biomarkers of response to azacitidine and nivolumab in relapsed/refractory acute myeloid leukemia: A nonrandomized, open-label, phase II study. Cancer Discov 9, 370–383 (2019).

11. Rutella, S., et al. Adaptive immune gene signatures correlate with response to flotetuzumab, a CD123 × CD3 bispecific DART^®^ molecule, in patients with relapsed/refractory acute myeloid leukemia. Blood 132, 444–444 (2018).

12. Chen, D.S. & Mellman, I. Elements of cancer immunity and the cancer-immune set point. Nature 541, 321–330 (2017).

13. Tumeh, P.C., et al. PD-1 blockade induces responses by inhibiting adaptive immune resistance. Nature 515, 568–571 (2014).

14. Zhao, J., et al. Immune and genomic correlates of response to anti-PD-1 immunotherapy in glioblastoma. Nat Med 25, 462–469 (2019).

15. Schalper, K.A., et al. Neoadjuvant nivolumab modifies the tumor immune microenvironment in resectable glioblastoma. Nat Med 25, 470–476 (2019).

16. Cloughesy, T.F., et al. Neoadjuvant anti-PD-1 immunotherapy promotes a survival benefit with intratumoral and systemic immune responses in recurrent glioblastoma. Nat Med 25, 477–486 (2019).

17. Taube, J.M., et al. Colocalization of inflammatory response with B7-H1 expression in human melanocytic lesions supports an adaptive resistance mechanism of immune escape. Sci Transl Med 4, 127ra137 (2012).

18. Ayers, M., et al. IFN-gamma-related mRNA profile predicts clinical response to PD-1 blockade. J Clin Invest 127, 2930–2940 (2017).

19. Ott, P.A., et al. T-cell-inflamed gene-expression profile, programmed death ligand 1 expression, and tumor mutational burden predict efficacy in patients treated with pembrolizumab across 20 cancers: KEYNOTE-028. J Clin Oncol 37, 318–327 (2019).

20. Benci, J.L., et al. Tumor interferon signaling regulates a multigenic resistance program to immune checkpoint blockade. Cell 167, 1540–1554 e1512 (2016).

21. Ng, S.W., et al. A 17-gene stemness score for rapid determination of risk in acute leukaemia. Nature 540, 433–437 (2016).

22. Danaher, P., Warren, S. & Cesano, A. Development of gene expression signatures characterizing the tumor-immune interaction. J Clin Oncol 36, 205–205 (2018).

23. Danaher, P., et al. Gene expression markers of Tumor Infiltrating Leukocytes. J Immunother Cancer 5, 18 (2017).

24. Mrozek, K., et al. Prognostic significance of the European LeukemiaNet standardized system for reporting cytogenetic and molecular alterations in adults with acute myeloid leukemia. J Clin Oncol 30, 4515–4523 (2012).

25. Vadakekolathu, J., et al. Immune gene expression profiling in children and adults with acute myeloid leukemia identifies distinct phenotypic patterns. Blood 130, 3942–3942 (2017).

26. Spranger, S., et al. Up-regulation of PD-L1, IDO, and Tregs in the melanoma tumor microenvironment is driven by CD8+ T cells. Sci Transl Med 5, 200ra116 (2013).

27. Blank, C.U., et al. Neoadjuvant versus adjuvant ipilimumab plus nivolumab in macroscopic stage III melanoma. Nat Med 24, 1655–1661 (2018).

28. Zemek, R.M., et al. Sensitization to immune checkpoint blockade through activation of a STAT1/NK axis in the tumor microenvironment. Sci Transl Med 11 (2019).

29. Ehninger, A., et al. Distribution and levels of cell surface expression of CD33 and CD123 in acute myeloid leukemia. Blood Cancer J 4, e218 (2014).

30. Radpour, R., et al. CD8(+) T cells expand stem and progenitor cells in favorable but not adverse risk acute myeloid leukemia. Leukemia (2019).

31. Bill, M., et al. A 17-gene leukemia stem cell (LSC) score in adult patients with acute myeloid leukemia (AML) reveals a distinct mutational landscape and refines current European Leukemianet (ELN) genetic risk stratification. Blood 132, 289–289 (2018).

32. Farrar, J.E., et al. Genomic profiling of pediatric acute myeloid leukemia reveals a changing mutational landscape from disease diagnosis to relapse. Cancer Res 76, 2197–2205 (2016).

33. Bolouri, H., et al. The molecular landscape of pediatric acute myeloid leukemia reveals recurrent structural alterations and age-specific mutational interactions. Nat Med 24, 103–112 (2018).

34. Majzner, R.G., Heitzeneder, S. & Mackall, C.L. Harnessing the immunotherapy revolution for the treatment of childhood cancers. Cancer Cell 31, 476–485 (2017).

35. Dolfi, D.V., et al. Increased T-bet is associated with senescence of influenza virus-specific CD8 T cells in aged humans. J Leukoc Biol 93, 825–836 (2013).

36. Paley, M.A., et al. Progenitor and terminal subsets of CD8+ T cells cooperate to contain chronic viral infection. Science 338, 1220–1225 (2012).

37. Walter, R.B., et al. Prediction of early death after induction therapy for newly diagnosed acute myeloid leukemia with pretreatment risk scores: a novel paradigm for treatment assignment. J Clin Oncol 29, 4417–4423 (2011).

38. Weichselbaum, R.R., et al. An interferon-related gene signature for DNA damage resistance is a predictive marker for chemotherapy and radiation for breast cancer. Proc Natl Acad Sci U S A 105, 18490–18495 (2008).

39. Tyner, J.W., et al. Functional genomic landscape of acute myeloid leukaemia. Nature 562, 526–531 (2018).

40. Valk, P.J., et al. Prognostically useful gene-expression profiles in acute myeloid leukemia. N Engl J Med 350, 1617–1628 (2004).

41. Wagner, S., et al. A parsimonious 3-gene signature predicts clinical outcomes in an acute myeloid leukemia multicohort study. Blood Adv 3, 1330–1346 (2019).

42. Bezzi, M., et al. Diverse genetic-driven immune landscapes dictate tumor progression through distinct mechanisms. Nat Med 24, 165–175 (2018).

43. Chichili, G.R., et al. A CD3xCD123 bispecific DART for redirecting host T cells to myelogenous leukemia: preclinical activity and safety in nonhuman primates. Sci Transl Med 7, 289ra282 (2015).

44. Uy, G.L., et al. Phase 1 cohort expansion of flotetuzumab, a CD123×CD3 bispecific Dart^®^ protein in patients with relapsed/refractory acute myeloid leukemia (AML). Blood 132, 764–764 (2018).

45. Al-Hussaini, M., et al. Targeting CD123 in acute myeloid leukemia using a T-cell-directed dual-affinity retargeting platform. Blood 127, 122–131 (2016).

46. Davidson-Moncada, J., Viboch, E., Church, S.E., Warren, S.E. & Rutella, S. Dissecting the immune landscape of acute myeloid leukemia. Biomedicines 6, 110 (2018).

47. Walter, R.B., et al. Resistance prediction in AML: analysis of 4601 patients from MRC/NCRI, HOVON/SAKK, SWOG and MD Anderson Cancer Center. Leukemia 29, 312–320 (2015).

48. Herold, T., et al. A 29-gene and cytogenetic score for the prediction of resistance to induction treatment in acute myeloid leukemia. Haematologica 103, 456–465 (2018).

49. Khodarev, N.N., et al. STAT1 is overexpressed in tumors selected for radioresistance and confers protection from radiation in transduced sensitive cells. Proc Natl Acad Sci U S A 101, 1714–1719 (2004).

50. Christopher, M.J., et al. Immune Escape of Relapsed AML Cells after Allogeneic Transplantation. N Engl J Med 379, 2330–2341 (2018).

51. Li, J., et al. Tumor cell-intrinsic factors underlie heterogeneity of immune cell infiltration and response to immunotherapy. Immunity 49, 178–193 e177 (2018).

52. Luke, J.J., Bao, R., Sweis, R.F., Spranger, S. & Gajewski, T.F. WNT/beta-catenin pathway activation correlates with immune exclusion across human cancers. Clin Cancer Res (2019).

53. Senbabaoglu, Y., et al. Tumor immune microenvironment characterization in clear cell renal cell carcinoma identifies prognostic and immunotherapeutically relevant messenger RNA signatures. Genome Biol 17, 231 (2016).

54. Dong, Z.Y., et al. Potential predictive value of TP53 and KRAS mutation status for response to PD-1 blockade immunotherapy in lung adenocarcinoma. Clin Cancer Res 23, 3012–3024 (2017).

55. Welch, J.S., et al. TP53 and decitabine in acute myeloid leukemia and myelodysplastic syndromes. N Engl J Med 375, 2023–2036 (2016).

## References

56. Sugio, T., et al. Microenvironmental immune cell signatures dictate clinical outcomes for PTCL-NOS. Blood Adv 2, 2242–2252 (2018).

57. Payton, J.E., et al. High throughput digital quantification of mRNA abundance in primary human acute myeloid leukemia samples. J Clin Invest 119, 1714–1726 (2009).

58. Subramanian, A., et al. Gene set enrichment analysis: a knowledge-based approach for interpreting genome-wide expression profiles. Proc Natl Acad Sci U S A 102, 15545–15550 (2005).

59. Rutella, S., et al. Capturing the complexity of the immune microenvironment of acute myeloid leukemia with 3D biology technology. J Clin Oncol 36, 50–50 (2018).

60. Stavropoulou, V., et al. MLL-AF9 expression in hematopoietic stem cells drives a highly invasive AML expressing EMT-related genes linked to poor outcome. Cancer Cell 30, 43–58 (2016).

61. Ley, T.J., et al. Genomic and epigenomic landscapes of adult de novo acute myeloid leukemia. N Engl J Med 368, 2059–2074 (2013).

62. Edgar, R., Domrachev, M. & Lash, A.E. Gene Expression Omnibus: NCBI gene expression and hybridization array data repository. Nucleic Acids Res 30, 207–210 (2002).

63. Lauten, M., et al. Prediction of outcome by early bone marrow response in childhood acute lymphoblastic leukemia treated in the ALL-BFM 95 trial: differential effects in precursor B-cell and T-cell leukemia. Haematologica 97, 1048–1056 (2012).

64. Metsalu, T. & Vilo, J. ClustVis: a web tool for visualizing clustering of multivariate data using Principal Component Analysis and heatmap. Nucleic Acids Res 43, W566–570 (2015).

