## Supplementary material for "Immune landscapes predict chemotherapy resistance and immunotherapy response in acute myeloid leukemia": Legends to supplemental figures

Extended Data Fig. 1: **A**) Pie chart showing patient distribution by European Leukemia Net (ELN) cytogenetic risk category in the AML discovery series (PMCC cohort; n=290 cases). NA = not available. **B**) Cohort-wide relapse-free survival (RFS) and overall survival (OS) in the AML discovery series (PMCC cohort; disease and patient characteristics are detailed in **Table 1**). The Kaplan-Meier method was used to generate survival curves, which were compared using a log-rank test (*p<0.05; ***p<0.001). Tick marks indicate censored observations. ELN=European Leukemia Net. **C**) Interferon (IFN), adaptive and myeloid signature scores individually separate AML cases according to high and low expression values. ClustVis, an online tool for clustering of multivariate data, was used for data analysis and visualization^1^. **D**) Violin plots summarizing the expression of leukemia-associated markers (CD33, CD34, CD123, CD117) in immune-infiltrated and immune-depleted AML cases from the PMCC cohort. Data were compared using the Mann-Whitney U test for unpaired determinations (*p<0.05; ***p<0.001; ns=not significant). **E**) Kaplan-Meier estimate of RFS in patients with AML separated by immune gene score quartiles (highest [n=78] and lowest [n=79]). Survival curves were compared using the log-rank test (*p<0.05; **p<0.01).

Extended Data Fig. 2: **A**) Unsupervised hierarchical clustering (Euclidean distance, complete linkage) of the correlation matrix of immuno-oncology proteins detected with GeoMx™ Digital Spatial Profiling (DSP) allows the identification of CD3 neighbors in 10 pre-treatment BM trephine biopsies from adult patients (SAL series) with high and low T-cell infiltration (median split). Correlation value color-coded *per* the legend. β2M=β2-microglobulin. Morpheus, an online tool developed at the Broad Institute (MA, USA) was used for data analysis and visualization. **B**) Unsupervised hierarchical clustering (Euclidean distance, complete linkage) of immune cell type-specific and biological activity scores (mRNA measurements) in 10 pre-treatment BM trephine biopsies used for GeoMx DSP. ClustVis, an online tool for clustering of multivariate data, was used for data analysis and visualization^1^. **C**) Correlation between CD3 protein expression, as determined by GeoMx DSP, and mRNA immune gene signatures (T-cell score, CD8 score, exhausted CD8 score, NK score, cytotoxicity score and macrophage score) as measured with the nCounter^®^ platform. **D**) Correlation between mRNA and protein expression, as determined by GeoMx DSP. Spearman rank correlation coefficients are shown in green; *p* values are shown in blue.

Extended Data Fig. 3: GeoMx™ digital spatial profiling (DSP) and region of interest (ROI) selection in a representative pre-treatment BM trephine biopsy (SAL series) with high levels of T-cell infiltration (CD3 shown in green). The median CD3 barcode count (24 ROIs) is indicated.

Extended Data Fig. 4: GeoMx™ digital spatial profiling (DSP) and region of interest (ROI) selection in a representative pre-treatment BM trephine biopsy (SAL series) with low levels of T-cell infiltration (CD3 shown in green). The median CD3 barcode count (24 ROIs) is indicated.

Extended Data Fig. 5: Correlation between immune gene signatures, which were derived as detailed in **Fig. 1**, and patients’ characteristics, including white blood cell (WBC) count (**A**), percentage of bone marrow (BM) blasts (**B**) and peripheral blood (PB) blasts (**C**), ELN cytogenetic risk category (**D**) and age at diagnosis (**E**) (PMCC series). Comparisons were performed using the Mann-Whitney *U* test for unpaired determinations. *P<0.05; ***P<0.0001.

Extended Data Fig. 6: **A-B**) Unsupervised hierarchical clustering (Euclidean distance, complete linkage) of the correlation matrix of immune and biological activity signatures highlights co-expression patterns of immune gene sets (correlation value color-coded *per* the legend; Pearson correlation coefficient >0.45; blue boxes) in bone marrow (BM) samples from children (CHOP cohort) and adults with AML (SAL cohort), namely, IFN-dominant signatures, adaptive immunity signatures and myeloid signatures. Morpheus, an online tool developed at the Broad Institute (MA, USA) was used for data analysis and visualization. **C-D**) IFN, adaptive and myeloid signature scores stratify children and adults with AML (CHOP and SAL series) into subgroups with high and low expression scores. ClustVis was used for data analysis and visualization^1^. **E**) IFN, adaptive and myeloid scores in aggregate stratify children and adults with AML (CHOP and SAL series) into subgroups with high and low expression scores. ClustVis was used for data analysis and visualization^1^. ELN=2017 European Leukemia Net. Criteria for response to induction chemotherapy (M1 through M3 in the top panel and early blast clearance [EBC]/complete remission [CR] in the bottom panel) are detailed in **Table 1**. ADV=adverse; INT=intermediate; FAV=favorable; NA=not available; NR=no response.

Extended Data Fig. 7: **A**) Top 20 differentially expressed (DE) immune genes between childhood and adult AML cases. ClustVis was used for data analysis and visualization^62^. Violin plots summarize the expression levels of relevant chemokine genes. Data were compared with the Mann-Whitney *U* test for paired determinations. Volcano plots of DE genes were generated using the nSolver^®^ software package. ***P<0.001. **B**) Top 20 DE immune genes between matched BM samples from adult patients (SAL cohort) at diagnosis and achievement of complete remission (CR). Data were compared with the Mann-Whitney *U* test for paired determinations. Volcano plots of DE genes were generated using the nSolver^®^ software package. ***P<0.001. **C**) Top 20 DE immune genes between matched BM samples from adult patients (SAL cohort) at diagnosis and disease relapse. Data were compared with the Mann-Whitney *U* test for paired determinations. Volcano plots of DE genes were generated using the nSolver^®^ software package. **P<0.01; ***P<0.001. **D**) Venn diagram showing minimal overlap in DE genes between children and adults with AML, and patients at disease onset, achievement of CR and relapse.

Extended Data Fig. 8: **A**) The Search Tool for the Retrieval of Interacting Genes/Proteins (STRING) database ([http://string-db.org](http://string-db.org/)) was used to critically assess and integrate protein-protein interactions between the top 20 genes differentially expressed in adults and children with AML, as well as across disease stages (diagnosis, achievement of complete remission [CR], relapse)^2^. A detailed list of significantly enriched Gene Ontology (GO) processes and KEGG pathways ranked by false discovery rate is provided as **Supplemental Tables 2-3**. Protein network analysis and predicted functional partners of DE genes with the highest-confidence interaction scores (>0.900) are shown. Network nodes (query proteins) represent proteins produced by a single, protein-coding gene locus. White nodes represent second shells of interactors. Empty and filled nodes indicate proteins of unknown or partly known 3D structure, respectively. Edges represent protein-protein associations. Line shapes denote predicted modes of action. **B**) GO enrichment analysis. For each given gene list submitted to metascape.org, pathway and process enrichment analyses were carried out using all genes in the genome as the enrichment background. Terms with a *p* value <0.01, a minimum count of 3, and an enrichment factor >1.5 (defined as the ratio between the observed counts and the counts expected by chance) were collected and grouped into clusters based on their membership similarities. **C**) Expression of genes associated with T-cell senescence/terminal differentiation/exhaustion (*TIGIT*, *TBX21* or T-bet, *CTLA4*)^3,4^ in matched BM samples from adult patients (SAL series) at disease onset and relapse (n=9). Data were compared with the Mann-Whitney *U* test for paired determinations. **P<0.01. ns=not significant.

Extended Data Fig. 9: **A**) Frequency, co-occurrence and mutual exclusivity of common molecular lesions with prognostic significance in AML cases with RNA sequencing data from The Cancer Genome Atlas (TCGA). Data were retrieved and analyzed using cBioPortal (<http://www.cbioportal.org/>). **B**) Locations and predicted consequences of *TP53* mutations identified in AML cases from TCGA.

Extended Data Fig. 10: Abnormalities of IFN downstream genes (missense mutations, mRNA upregulation, amplification and deep deletion) in relation to *TP53* mutations in AML cases from TCGA. Data were retrieved and analyzed using cBioPortal (<http://www.cbioportal.org/>).

Extended Data Fig. 11: Immune subtypes associate with cancer driver gene mutations in The Cancer Genome Atlas (TCGA)-AML and Beat AML trial specimens. **A**) Unsupervised hierarchical clustering (Euclidean distance, complete linkage) of immune cell type signatures and biological activity signatures in TCGA-AML cases stratified by prognostic molecular lesions. ClustVis was used for data analysis and visualization^1^. Blue arrows denote *TP53* mutated cases; green arrows identify *RUNX1*-mutated cases. **B**) Expression of IFN-related gene sets (as defined in **Fig. 1**) in TCGA cases with *TP53* mutations (adverse molecular risk), *NPM1* mutations (ELN favorable risk), *RUNX1* mutations (adverse molecular risk), *NPM1* mutations with *FLT3*-ITD (ELN intermediate risk), FLT3-*ITD* alone (ELN adverse risk) or other mutations (*TET2*, *DNMT3A*, *IDH1*, *IDH2*). Comparisons were performed using the Kruskal-Wallis test for unpaired determinations. **C**) Expression of IFN-downstream genes in Beat AML cases with (n=21) or without *TP53* mutations (n=128). ClustVis was used for data analysis and visualization^1^. **D**) Violin plots summarizing the expression of surrogate markers of T-cell infiltration and cytotoxicity in Beat AML cases with or without *TP53* mutations. Comparisons were performed with the Mann-Whitney *U* test for unpaired data. *P<0.05; **P<0.01.

**References**

1.	Metsalu, T. & Vilo, J. ClustVis: a web tool for visualizing clustering of multivariate data using Principal Component Analysis and heatmap. *Nucleic Acids Res* **43**, W566-570 (2015).

2.	Szklarczyk, D.*, et al.* The STRING database in 2011: functional interaction networks of proteins, globally integrated and scored. *Nucleic Acids Res* **39**, D561-568 (2011).

3.	Paley, M.A.*, et al.* Progenitor and terminal subsets of CD8+ T cells cooperate to contain chronic viral infection. *Science* **338**, 1220-1225 (2012).

4.	Dolfi, D.V.*, et al.* Increased T-bet is associated with senescence of influenza virus-specific CD8 T cells in aged humans. *J Leukoc Biol* **93**, 825-836 (2013).
